# Histone H3 serotonylation dynamics in dorsal raphe nucleus contribute to stress- and antidepressant-mediated gene expression and behavior

**DOI:** 10.1101/2023.05.04.539464

**Authors:** Amni Al-Kachak, Sasha L. Fulton, Giuseppina Di Salvo, Jennifer C Chan, Lorna A. Farrelly, Ashley E. Lepack, Ryan M. Bastle, Lingchun Kong, Flurin Cathomas, Emily L. Newman, Caroline Menard, Aarthi Ramakrishnan, Polina Safovich, Yang Lyu, Herbert E. Covington, Li Shen, Kelly Gleason, Carol A. Tamminga, Scott J. Russo, Ian Maze

**Affiliations:** Nash Family Department of Neuroscience, Friedman Brain Institute, Icahn School of Medicine at Mount Sinai, New York, New York 10029, USA; Department of Psychiatry, McLean Hospital and Harvard Medical School, Belmont, MA 02478, USA; Department of Psychology, Empire State College, State University of New York, Saratoga Springs, NY 12866; Department of Psychiatry, University of Texas Southwestern Medical School, Dallas, TX, 75390, USA; Department of Pharmacological Sciences, Icahn School of Medicine at Mount Sinai, New York, New York 10029, USA; Howard Hughes Medical Institute, Icahn School of Medicine at Mount Sinai, New York, New York 10029, USA

**Author notes:** **Corresponding author:** Ian Maze.

**Keywords:** Chronic social defeat stress, dorsal raphe nucleus, histone serotonylation, antidepressants, ChIP-seq, RNA-seq

## Abstract

**Background:** Major depressive disorder (MDD), along with related mood disorders, is a debilitating illness that affects millions of individuals worldwide. While chronic stress increases incidence levels of mood disorders, stress-mediated disruptions in brain function that precipitate these illnesses remain elusive. Serotonin-associated antidepressants (ADs) remain the first line of therapy for many with depressive symptoms, yet low remission rates and delays between treatment and symptomatic alleviation have prompted skepticism regarding precise roles for serotonin in the precipitation of mood disorders. Our group recently demonstrated that serotonin epigenetically modifies histone proteins (H3K4me3Q5ser) to regulate transcriptional permissiveness in brain. However, this phenomenon has not yet been explored following stress and/or AD exposures.

**Methods:** We employed a combination of genome-wide and biochemical analyses in dorsal raphe nucleus (DRN) of male and female mice exposed to chronic social defeat stress to examine the impact of stress exposures on H3K4me3Q5ser dynamics, as well as associations between the mark and stress-induced gene expression. We additionally assessed stress-induced regulation of H3K4me3Q5ser following AD exposures, and employed viral-mediated gene therapy to reduce H3K4me3Q5ser levels in DRN and examine the impact on stress-associated gene expression and behavior.

**Results:** We found that H3K4me3Q5ser plays important roles in stress-mediated transcriptional plasticity. Chronically stressed mice displayed dysregulated H3K4me3Q5ser dynamics in DRN, with both AD- and viral-mediated disruption of these dynamics proving sufficient to rescue stress-mediated gene expression and behavior.

**Conclusions:** These findings establish a neurotransmission-independent role for serotonin in stress-/AD-associated transcriptional and behavioral plasticity in DRN.

## INTRODUCTION

Major depressive disorder (MDD), along with related mood disorders, is an enigmatic and highly heterogeneous syndrome that affects approximately 17 million American adults each year (1). Chronic stress exposures represent a major risk factor for MDD (2), however the molecular mechanisms underlying stress-induced susceptibility to depression remain poorly understood. Despite being serendipitously discovered more than 60 years ago, antidepressant (AD) treatments that target monoaminergic systems (e.g., selective serotonin reuptake inhibitors/SSRIs) remain the first line of therapy for many with MDD. Yet long delays between initiation of treatment and symptomatic alleviation, along with low remission rates (3), have encouraged further investigation to identify more direct therapeutic targets. The monoamine neurotransmitter serotonin, or 5-hydroxytryptamine (5-HT), in particular, is thought to play critical roles in neuronal plasticity associated with affective disorders, as altered serotonergic signaling is implicated in both the etiology and treatment of MDD (4). However, a recent report revealing a lack of robust evidence linking alterations in serotonin levels, MDD and AD efficacy has prompted renewed interest from the field in defining precise roles for 5-HT in the precipitation and treatment of MDD (5).

In the central nervous system, 5-HT has long been thought to function primarily as a neuromodulator, regulating a wide array of physiological and behavioral functions, including cognitive and emotional processing, autonomic control and sleep-wake cycles (6). In the brain, 5-HT is synthesized predominantly in monoaminergic, tryptophan hydroxylase 2-expressing neurons located in the dorsal raphe nucleus (DRN). 5-HT is thought to elicit its neuromodulatory effects via a complex and wide-ranging efferent system that projects broadly throughout the brain (including to key regions of the limbic system, such as the prefrontal cortex, nucleus accumbens and amygdala, as well as the hippocampus and cerebellum, among others) (7). In this well-documented view, 5-HT receptor-mediated mechanisms initiate alterations in cell-cell communication, which in turn can contribute to the plasticity of postsynaptic neurons (8–10). During early brain development, 5-HT can additionally act as a trophic factor to regulate neuronal growth and differentiation processes, synaptogenesis and dendritic pruning (11–13), suggesting potential roles for this molecule beyond its actions as a neurotransmitter. Along these lines, while SSRIs function pharmacologically to perturb 5-HT signaling in brain via inhibition of the 5-HT transporter (SLC6A4/SERT) – a phenomenon that contributed to the development of the ‘monoamine hypothesis of depression’ (14) – it remains unclear whether serotonergic dysfunction itself promotes MDD-related pathologies, or how therapeutics might work mechanistically to promote symptomatic alleviation in MDD individuals.

While vesicular packaging of monoamines is essential for neurotransmission, previous data demonstrated the additional presence of extravesicular monoamines in both the soma and nucleus of monoaminergic neurons (15,16). In addition to its role as a neuromodulator, 5-HT was previously shown to be capable of forming covalent bonds with certain substrate proteins via transamidation by the tissue Transglutaminase 2 enzyme, a process referred to as serotonylation (17). In more recent studies, our group identified a new class of histone posttranslational modification (PTM) termed monoaminylation, whereby monoamine neurotransmitters, such as 5-HT, dopamine and histamine, can be transamidated onto histone glutamine residues (18–23). We showed that histone H3 glutamine (Q) 5 is a primary site for these PTMs and demonstrated that H3 monoaminylation states play important roles in the regulation of neuronal transcriptional programs, both during early development/cellular differentiation and in adult brain. We demonstrated that combinatorial H3 lysine 4 (K4) tri-methylation (me3) glutamine 5 (Q5) serotonylation (H3K4me3Q5ser), in particular, acts as a permissive epigenetic mark, both by enhancing the binding of the general transcription factor complex TFIID, and attenuating H3K4me3 demethylation via inhibition of K4me3 demethylases (18,24). While these PTMs play critical roles in the regulation of normal patterns of transcription in brain, we also found that certain H3 monoaminylations (e.g., H3 dopaminylation) are inappropriately dynamic in response to aberrant environmental stimuli, which contribute to maladaptive neuronal plasticity in disorders associated with altered monoaminergic signaling (e.g., cocaine and opiate use disorders) (19,25,26).

Given the chronic, relapsing nature of MDD, great efforts have been taken over the past two decades to examine the underlying molecular determinants of this brain disorder, the findings of which have uncovered various patterns of transcriptional and epigenetic dysregulation – often brain region and cell-type specific – as potential causative factors in the precipitation and persistence of MDD-related pathophysiology (27,28). Furthermore, explorations in preclinical rodent models of chronic stress, which can be used to model specific endophenotypes associated with MDD (e.g., anhedonia, social avoidance, behavioral despair, cognitive deficits, etc.), have revealed strong correlations between epigenetic dysfunction, gene expression abnormalities and behavioral stress susceptibility (29–34). However, our understanding of how these mechanisms mediate life-long susceptibility to stress-induced syndromes like MDD remains limited. Additionally, while much evidence exists implicating molecular alterations in cortical and limbic brain structures (all of which receive dense serotonergic projections) as precipitating factors in the regulation of stress susceptibility *vs.* resilience (35–40), fewer studies have explored chromatin-related phenomena in DRN, which may also contribute significantly to behavioral dysregulation in affect-related disorders.

Here, we demonstrate that DRN displays significant alterations in mood disorder-associated gene expression programs following chronic social stress in both male and female mice, with behaviorally resilient *vs.* susceptible animals displaying blunted transcriptional abnormalities. We further show that histone H3 serotonylation patterns are reorganized in response to chronic stress in both sexes, a phenomenon that is rescued in both behaviorally resilient animals and mice chronically treated with the SSRI fluoxetine. Finally, we demonstrate that directly reducing levels of H3 serotonylation in DRN using a dominant negative viral vector approach is sufficient to reverse chronic stress-induced gene expression programs and promote behavioral resilience to stressful stimuli. In sum, these findings establish a non-canonical, neurotransmission-independent role for 5-HT in stress-mediated transcriptional and behavioral plasticity in DRN, and indicate that certain ADs may function, at least in part, to reverse altered patterns of H3 serotonylation in brain.

## MATERIALS AND METHODS

### ANIMALS

C57BL/6J mice were purchased from The Jackson Laboratory. Retired male CD-1 breeders of at least 4 months of age were purchased from Charles River laboratories and used as aggressors. All mice were singly housed following CSDS and maintained on a 12-h/12-h light/dark cycle throughout the entirety of the experiments. Mice were provided with *ad libitum* access to water and food throughout the entirety of the experiments. All animal procedures were done in accordance with NIH guidelines and with approval with the Institutional Animal Care and Use Committee of the Icahn School of Medicine at Mount Sinai.

### MALE CSDS

Male chronic social defeat stress (CSDS) was performed, as previously described (36). Briefly, CD-1 retired breeders were screened for aggressive behavior and were then single-housed in static hamster cages on one side of a clear perforated divider 24 hr prior to the start of CSDS. For 10 min every day, for 10 days, 8-week old C57BL/6J experimental mice were placed in the same side of the home cage as the CD-1 mouse. The CD-1 mouse was then allowed to physically attack the intruder C57BL/6J mouse throughout the 10-min defeat session. After each defeat session, experimental mice were moved to the opposite side of the clear perforated divider for 24 hr, permitting sensory interactions with the aggressor. Experimental mice were then rotated to a new cage with a novel aggressor every day for the remainder of the experiments. 24 hr after the final defeat, experimental mice were single-housed in static mouse cages for subsequent social interaction testing.

*Controls:* 8-week old C57BL/6J control mice were pair-housed in mouse cages on either side of a clear perforated divider, similar to the ones used in hamster cages. Each control mouse was exposed to a novel mouse daily via rotation in a similar fashion to the experimental animal, but was never exposed to a CD-1 aggressor. Control mice were single-housed in static mouse cages at the end of the 10-d experiment for subsequent social interaction testing.

All behavioral protocols adhered strictly to the Guidelines for the Care and Use of Mammals in Neuroscience and Behavioral Research (National Academies Press, Washington, DC, 2003). All animals subjected to any form of stress were carefully monitored for their health and wellbeing in concert with the Icahn School of Medicine at Mount Sinai’s veterinary staff. Any animals showing untoward effects of stress were euthanized. In our experience, such untoward effects are extremely rare (<3% of all animals studied).

### MALE SI TESTING

24 hr after completion of CSDS, mice were tested for social avoidance via social interaction testing, as described previously (36). In this test, animals were transferred to a quiet room under red-light conditions and were habituated for 30 min to 1 hr prior to testing. For the first session, the subject animal was placed in a novel open-field arena with a small, wired enclosure on one side of the arena. The mouse was allowed 2.5 min to explore the empty arena, and its baseline exploration behavior was tracked from above via a video camera connected to a computer running Ethovision tracking software. In the second session, a novel CD-1 mouse was placed in the small enclosure in the arena, and the subject mouse was placed back in the arena for another 2.5 min, and exploration behavior was tracked via EthoVision. Social interaction was assessed by SI ratio, which is the amount of time the animal spent in the interaction zone while the CD-1 mouse was present, over the time spent in the interaction zone while the CD-1 was absent. A subject mouse was deemed to be stress-resilient if it had an SI ratio greater than 1, whereas stress-susceptible mouse had SI ratios less than 1.

### FEMALE CSDS

Female social defeat was performed as previously described (41). Briefly, intact female Swiss Webster (CFW) mice were housed with castrated male mice and were tested for aggression against experimental female intruder mice. Wild-type 12-week old female C57BL/6J (B6) mice were socially defeated daily by aggressive CFW female resident mice for 5 minutes per day during the 10-day paradigm. Between the defeats, experimental B6 female mice were housed with the aggressor female in a shared home cage, separated by a clear perforated cage divider. Control females were housed in identical conditions but were never exposed to a physical defeat. Defeated and control females were singe housed following the final defeat.

### FEMALE SI TESTING

Social interaction testing was done in the experimental female’s home cage 24 hours after the final defeat. In this test, a non-aggressive B6 female was placed into the experimental female’s home cage for 1.5 minutes and social interaction time and defensive score was assessed. Social interaction included any anogenital, flank, naso-nasal sniffing, or flank on flank contact that was initiated by the experimental animal. Defensive score was defined on a numerical scale from 0-3, with 0 being not defensive, 1 being minimally defensive (avoidance only), 2 being moderately defensive (avoidance, digging, but no kicking), and 3 being highly defensive (avoidance, escape, kicking, flinching, digging, jumping, pacing). Tissue was collected 24 hours after the social interaction test (i.e., 48 hours after the final defeat). As in previous reports using this female CSDS paradigm (41), vaginal cytology was monitored in experimental mice during the 10-day social defeat protocol using the lavage technique. Consistent with the literature, CSDS did not affect estrous cycling (nor body weight) in defeated females. Since we did not have any evidence to suggest that estrous stage significantly impacts female responsiveness to CSDS, we did not use it as a covariate in our sequencing analyses.

### FLUOXETINE TREATMENTS

24 hr following social interaction testing, each group of male mice (control, stress-susceptible and stress-resilient) were randomly separated into two groups, either to receive regular drinking water (vehicle) or drinking water with fluoxetine hydrochloride for 30 d. Drug treatment was performed as previously described (42). Briefly, fluoxetine hydrochloride (Spectrum Chemical) was administered *ad libitum* in drinking water (filtered tap water) in opaque light-protected bottles (Argos Technologies Litesafe Centrifuge Tubes 50mL-Fisher, #03-395-120). Fluoxetine solutions were changed and refreshed 3 times per week. Fluoxetine was administered through drinking water at 160 mg/L. Water was weighed every day to monitor consumption and track dosage. Mice drank ∼ 2–3 ml per day of 160 mg/L solution, resulting in an estimated 15.25 mg/kg dose over the treatment period. Following completion of 30 d of treatment, mice underwent social interaction testing to evaluate drug efficacy.

### FORCED SWIM TEST (FST)

The forced swim test was conducted as previously described (43). Mice were placed in a 4-liter glass beaker with 2L of room-temperature water for 6 minutes. Each session was recorded and hand-scored, recording the number of seconds the mouse was immobile.

### OPEN FIELD TEST (OFT)

Open field testing was performed as previously described (43). Mice were placed in a 16 x 16-inch open field apparatus under dim lighting and distance and time in center *vs.* periphery were recorded via Ethovision software.

### ELEVATED PLUS MAZE (EPM)

The elevated plus maze was used as previously described (43). Mice were placed into the center of the maze under dim lighting and allowed to explore for 5 min. Time spent in the closed and open arms and number of explorations of open arms was recorded via Ethovision software, as previously described (43).

### HUMAN BRAIN SAMPLES

Human DRN tissues from the Dallas Brain Collection (UT Neuropsychiatry Research Program) were obtained from the Southwestern Institute of Forensic Sciences at Dallas, UT Southwestern Transplant Services Center, and UT Southwestern Willed Body Program, following consent from donor subjects’ next of kin, permission to access medical records and to hold direct telephone interviews with a primary caregivers. All clinical information obtained for each donor was reviewed by three research psychiatrists, using DSM-V criteria for diagnoses. Blood toxicology screens were conducted for each donor subject from the Southwestern Institute of Forensic Sciences at Dallas. Collection of postmortem human brain tissues is approved by the University of Texas Southwestern Medical Center Institutional Review Board [STU 102010-053]. Brain tissue dissections were removed, frozen immediately using dry ice and 2-methylbutane (1:1, v:v) and stored at –80 °C. For western blotting validation experiments, H3 was used an internal reference control for the best normalization and most reliable indicator of equal protein concentration. Demographic information can be found in **Supplemental Data Table 70**.

### RNA ISOLATION, RNA-SEQ AND ANALYSIS

For male and female CSDS experiments, DRN tissues were collected from mice 24 hr after final social interaction (1 mm punches) and immediately flash-frozen. To examine genome-wide effects of blocking serotonylation via viral infection, brains were sectioned at 100 µm on a cryostat, and GFP/RFP was illuminated with a NIGHTSEA BlueStar flashlight to microdissect virally infected tissues. DRN tissue punches were homogenized in Trizol (Thermo Fisher), and RNA was isolated on RNeasy Microcolumns (Qiagen) following manufacturer’s instructions. Following RNA purification, RNA-seq libraries were prepared according to the Illumina Truseq RNA Library Prep Kit V2 (#RS-122-2001) protocol and sequenced on the Illumina Novaseq platform. Following sequencing, data was pre-processed and analyzed as previously described (19). Briefly, FastQC (Version 0.72) was performed on the concatenated replicate raw sequencing paired-end reads from each library to ensure minimal PCR duplication and sequencing quality. Reads were aligned to the mouse mm10 genome using HISAT2 (Version 2.1.0) and annotated against Ensembl v90. After removal of multiple-aligned reads, remaining reads were counted using featurecounts (Version 2.0.1) with default parameters, and filtered to remove genes with low counts (<10 reads across samples). For male 24-hr post-CSDS RNA-seq, RUVr (44), k = 6, was performed to normalize read counts based on the residuals from a first-pass GLM regression of the unnormalized counts on the covariates of interest. For female RNA-seq experiments and the serotonylation manipulation experiments with the Q5A virus, RUVr (44) (female; k = 4, Q5A; k = 6 was performed to normalized read counts. DESEQ2 (45) (Version 2.11.40.6) was used to perform pairwise differential expression analyses between indicated comparisons. Differentially expressed (DE) genes were defined at FDR<0.05. Unsupervised clustering heatmaps were generated at DE genes across samples using heatmap2 from gplots (Version 3.1.3). Threshold free Rank-Rank Hypergeometric Overlap (RRHO) maps were generated to visualize transcriptome-wide gene expression concordance patterns as previously described (46), using RRHO2 (Version 1.0). Odds ratios for overlapping gene sets were calculated with GeneOverlap (Version 1.34.0). Enrichment analysis on gene sets of interest was performed with EnrichR, Benjamini-Hochberg (BH) q-values corrected for multiple testing are reported (47–49).

### WESTERN BLOTTING AND ANTIBODIES

DRN tissues were collected from mice (1 mm punches) and immediately flash-frozen. Punches were homogenized using a sonicator in RIPA Buffer, containing 50 mM Tris-HCl, 150 mM NaCl, 0.1% SDS, 1% NP-40 and 1x protease inhibitor cocktail. Protein concentrations were measured using the DC protein assay kit (BioRad), and 20 ug of protein was loaded onto 4-12% NuPage BisTris gels (Invitrogen) for electrophoresis. Proteins were then fast-transferred using nitrocellulose membranes and blocked for 1 hr in 0.1% Tween-20 in 1x PBS (PBS-T) in a 5% milk buffer, before undergoing overnight incubation with primary antibodies at 4° C. The following day, blots were washed of primary antibody for 10 min 3X in PBS-T, then incubated for 1 hr with horseradish peroxidase conjugated anti-rabbit (BioRad 170-6515, lot #: 64033820) or anti-mouse (GE Healthcare UK Limited NA931V, lot #: 9814763) secondary antibodies (1:10000; 1:50000 for anti-H3 antibody, BioRad) in 0.1% Tween-20 in 1x PBS (PBS-T) in a 5% milk buffer at RT. Blots were then washed of secondary antibody for 10 min 3X in PBS-T and bands were detected using enhanced chemiluminescence (ECL; Millipore). Densitometry was used to quantify protein bands via Image J Software and proteins were normalized to total H3 or GAPDH, as indicated. For cultured cerebellar granule neuron (cGN) western blotting experiments, 1 hour after 50 mM KCl treatment, cGNs in 6-well plates were rinsed with 1x PBS and lysed in 200 ul of 2x SDS loading buffer (100 mM Tris-HCl pH 6.8, 20% glycerol, 4% SDS, 0.1% bromophenol blue and 2% 2-mercaptoethanol). 15 ul of samples were loaded on 4–12% NuPAGE gel and transferred to nitrocellulose membranes. The following antibodies were used: rabbit anti-H3K4me3Q5ser (1:500, ABE2580; MilliporeSigma), rabbit anti-H3Q5ser (1:500, MilliporeSigma; ABE1791,), rabbit anti-H3 (1:50000, Abcam ab1791), H4 (1:10000, Abcam; ab10158), H3.3 (1:2000, MilliporeSigma; 09-838,), FLAG (1:5000, Sigma; F3165,) and rabbit anti-Gapdh (1:10000, Abcam; ab9485).

### CHROMATIN IMMUNOPRECIPITATION

DRN tissues were collected from mice (1 mm punches) and immediately flash-frozen. Punches were crosslinked with 1% formaldehyde and rotated gently at room temperature for 12 minutes. Punches were then immediately quenched with glycine and rotated gently at room temperature for 5 minutes. Samples were washed thoroughly before lysis and sonications were performed, as previously described (18). Samples were then incubated with specific antibodies (7.5 μg per sample) bound to M-280 Dynabeads on a rotator at 4 °C overnight. The following day, immunoprecipitates were washed, eluted and reverse-crosslinked. Samples underwent RNA and protein digestion and DNA was purified using a Qiagen PCR purification kit. The following antibodies were used: rabbit anti-H3K4me3Q5ser (1:500, ABE2580; MilliporeSigma).

### CHIP-SEQ LIBRARY PREPARATION AND ANALYSIS

Following DNA purifications, ChIP-seq libraries were generated according to Illumina protocols and sequenced on an Illumina HiSeq2500, 4000 or Novaseq Sequencers. ChIP-seq peaks were called and differential analysis conducted exactly as described previously (18,50). Briefly, raw sequencing reads were aligned to the mouse or human genome (mm10 or hg38, respectively) using default settings of HISAT2. Alignments were filtered to only include uniquely mapped reads using SAMtools v.1.8. Peak-calling was normalized to respective inputs for each sample and was performed using MACS v.2.1.1 (51) with default settings and filtered for FDR< 0.05. Differential analysis was performed using diffReps (52) with a 1 kb window size. Peaks and differential sites were further annotated to nearby genes or intergenic regions using the region analysis tool from the diffReps package. To be considered a “real” peak-containing PCG, a significant peak (FDR<0.05, >5-fold enrichment over input) had to be found in a PCG (promoter and/or gene body) in at least one of: 3 conditions for male 24 hr post-SI testing (Control, Susceptible or Resilient); 2 conditions for female 24 hr post-SI testing (Control, Defeat); 4 conditions for fluoxetine experiments 30d post-SI testing (control −/+ FLX, SUS −/+ FLX); or 4 conditions for human DRN (MDD – ADs, MDD + ADs *vs.* matched controls). To be considered a differentially enriched gene, it had to first pass the aforementioned criteria, and then display a ≥1.5 or ≤ –1.5 or ≥1.0 or ≤ –1.0 fold difference between conditions (pairwise comparisons) at FDR<0.05 (as indicated throughout). Enrichment analysis on gene sets of interest was performed with EnrichR, Benjamini-Hochberg (BH) q-values corrected for multiple testing are reported (47–49).

## CHIP/RE-CHIP EXPERIMENTS IN CULTURED GRANULE NEURONS

### Cerebellar granule neuron culture

Granule neurons were prepared from cerebellum of P7 CD-1 mouse pups as previously described (53). On day 1 *in vitro* (*DIV 1*), granule neurons were transduced with AAV-empty, AAV-H3.3-WT or AAV-H3.3Q5A respectively. 2 days after infection, the medium was changed to low KCl medium (Basal Medium Eagle, GIBCO+5% Hyclone bovine growth serum, Cytiva+1x penicillin-streptomycin, GIBCO+1x GlutaMAX™ Supplement, GIBCO+5 mM KCl).

### ChIP and Re-ChIP-qPCR

ChIP assays were performed with cultured granule neurons, as described previously with modifications (54). After quenching and sonication, 15 million granule neurons and 15 ul anti-FLAG beads (Sigma, #A2220) were used for each ChIP reaction. After IP, chromatin was eluted twice with 100 ul of 3X FLAG Peptide solution (Sigma, #F4799, dissolved in ChIP lysis buffer) for 30 minutes at 4 °C. The two eluents were mixed and incubated with 1 ug of anti-H3K4me3Q5ser antibody (Millipore, #ABE2580) for 4h-overnight. Next, the chromatin-antibody mixture was incubated with 25 ul washed Dynabeads Protein A (Invitrogen, #10001D). The following steps were the same as the ChIP assays described above. ChIP and Re-ChIP DNA was purified using a Qiagen PCR purification kit and eluted in 60 ul elution buffer. 2 ul of ChIP or Re-ChIP DNA was used for each qPCR reaction. See **Supplemental Data Table 71** for mouse ChIP-qPCR primers.

### VIRAL CONSTRUCTS

Lentiviral constructs were generated as previously described (18). Briefly, lenti-H3.3 constructs [wildtype (WT) vs. (Q5A)-Flag-HA] were cloned into a pCDH-RFP vector via PCR and enzyme restriction digestion. Plasmids were purified and sent to GENEWIZ for sequence validation. pCDH-GFP-H3.3 plasmids were then sent to Cyagen Biosciences for lentiviral packaging. For cultured cerebellar granule neuron experiments, pAAV-CMV-H3.3-IRES-GFP constructs [wildtype (WT) *vs.* (Q5A)-Flag-HA *vs.* empty] were packaged as follows: 70%-80% confluent HEK293T cells were transfected pAAV2/1 (Addgene 112862), pAdDeltaF6 (Addgene 112867), and pAAV-CMV-IRES-GFP or pAAV-CMV-H3.3-WT-IRES-GFP or pAAV-CMV-H3.3-Q5A-IRES-GFP with PEI reagent (Polysciences, #26008-5). 48-72 hours after transfection, the media with AAVs were collected by centrifuge. AAV particles were precipitated by adding 10% volume of PEG 8000-NaCl solution (40% PEG 8000, 2.5 M NaCl, pH 7.4). Next, the AAV particles were resuspended in granule neurons culture medium.

### VIRAL TRANSDUCTION

Mice were anesthetized with a ketamine/xylazine solution (10/1 mg/kg) i.p. and positioned in a stereotaxic frame (Kopf instruments). 1 µl of viral construct was infused intra-DRN using the following coordinates; anterior-posterior (AP) −4.40mm, medial-lateral (ML) 0.0mm, dorsal-ventral (DV) −3.40mm. Following surgery, mice received meloxicam (1 mg/kg) s.c. and topical antibiotic treatments for 3 days. Chronic social defeat stress and other behaviors were performed at least 21 days post-surgery to allow for optimal viral expression and recovery.

### IMMUNOHISTOCHEMISTRY

Mice were anesthetized with ketamine/xylazine (10/1 mg/kg) i.p., and then perfused with cold phosphate buffered saline (PBS 1X) and 4% paraformaldehyde (PFA) in 1X PBS. Whole brains were then post-fixed in 4% PFA overnight at 4° C and then transferred into a solution of 30% sucrose/PBS 1X for two days. Brains were sliced into serial 40 µm coronal slices using a Leica (type) cryostat. Free-floating DRN slices were washed for 10 min 3X in PBS 1X, then incubated for 30 min in 0.2% Triton X/PBS 1X, then incubated for 1 hr at RT in blocking buffer (0.3% Triton X, 3% donkey serum, 1X PBS). Finally, slices were incubated overnight at 4° C with mouse anti-RFP (1:200; lot#: GR3181906-1, Abcam ab65856) and HA-488 (1:200; lot #: K1716, Life Technologies, Alexa Fluor SC-805). The following day, slices were washed for 10 min 3X in 1x PBS and then incubated for 2 hr at RT with a fluorescent-tagged Alexa Fluor 568 anti-mouse secondary antibody (1:500, lot #: 1218263, Life Technologies A11004). Slices were then washed for 10 min 3X in 1x PBS and incubated with DAPI (1:10000, lot #: RK2297251, Thermo Scientific 62248) for 10 min at RT before being mounted on Superfrost Plus slides (Fischer Scientific) and coverslipped with Prolong Gold (Invitrogen). Immunofluorescence was visualized using a confocal microscope (Zeiss LSM 780).

### STATISTICS

Statistical analyses were performed using Prism GraphPad software. For all behavioral testing and biochemical experiments involving more than two conditions, two-way or one-way ANOVAs were performed with subsequent *post hoc* analyses. For experiments comparing only two conditions, two-tailed Student’s t tests were performed. Sequencing-based statistical analyses are described above. In biochemical and RNA-seq analyses, all animals used were included as separate *n*s (i.e., samples were not pooled). In ChIP-seq analyses, animals were pooled per *n* as designated above. Significance was determined at p≤0.05. Where applicable, outliers were determined using the Grubb’s test (alpha = 0.05; noted in **Fig. S7**). All bar/dot plot data are represented as mean ± SEM.

## RESULTS

### Gene expression programs in DRN are responsive to chronic social stress

To begin investigating the impact of chronic stress exposures on gene expression programs in DRN, we performed chronic social defeat stress (CSDS) in adult male mice, a well-characterized and etiologically relevant rodent model for the study of human depression, which recapitulates numerous pathophysiological features of MDD (e.g., social avoidance, anhedonia, stress-related metabolic syndromes, etc.) and displays symptomatic reversal in response to chronic, but not acute, AD treatments (36,55,56). CSDS in male mice produced two distinct groups of stress-susceptible *vs.* stress-resilient animals, with stress-susceptible mice displaying heightened levels of social avoidance in comparison to resilient and control (i.e., non-stressed, handled) groups (**Fig. 1A**). We then performed bulk RNA-seq on DRN tissues from control *vs.* susceptible *vs.* resilient mice, followed by differential expression analysis to compare the three groups. We found that stress-susceptible male mice exhibited significant alterations in the expression of 2,266 protein-coding genes (PCGs; FDR<0.1) *vs.* respective handled controls. Subsequent unsupervised clustering of all three groups at these differentially regulated transcripts revealed a clear pattern of separation between stress-susceptible *vs.* control animals, with resilient animals displaying a pattern more similar to that of controls (only 56 PCGs were found to be differentially regulated comparing resilient *vs.* control mice; FDR<0.1) (**Fig. 1B, Supplemental Data Tables 1, 8-9**). Subsequent gene set enrichment analyses on dysregulated loci observed in susceptible *vs.* control mice [GO Biological Process and DisGeNET, the latter of which curates a collection of genes and variants associated with human diseases, integrating data from publicly available genomics repositories, GWAS catalog, animals models (focused on genotype x phenotype relationships) and current scientific literature] identified significant (FDR<0.05) overlaps with pathways/processes involved in neuronal development (e.g., axon guidance, axonogenesis, regulation of cell migration, etc.) and synaptic transmission (e.g., chemical synaptic transmission), as well as enrichment in disease associated pathways related to psychiatric and affect-related disorders (e.g., MDD and schizophrenia) (**Fig. 1C, Supplemental Data Tables 2-3**). Interestingly, while both up-(988) and downregulated (1,278) gene expression was observed in stress-susceptible male DRN *vs.* controls, downregulated genes appear to have contributed more significantly to gene ontology and disease pathway enrichment observed in **Fig. 1C** (FDR<0.05; **Data Fig. S1A-B, Supplemental Data Tables 4-7**). These findings suggest that stress-susceptible gene expression programs in male DRN, particularly those genes that are acutely repressed in response to chronic stress, may be relevant to aberrant patterns of neuronal and/or behavioral plasticity observed in response to chronic stress exposures.

**Fig. 1.**
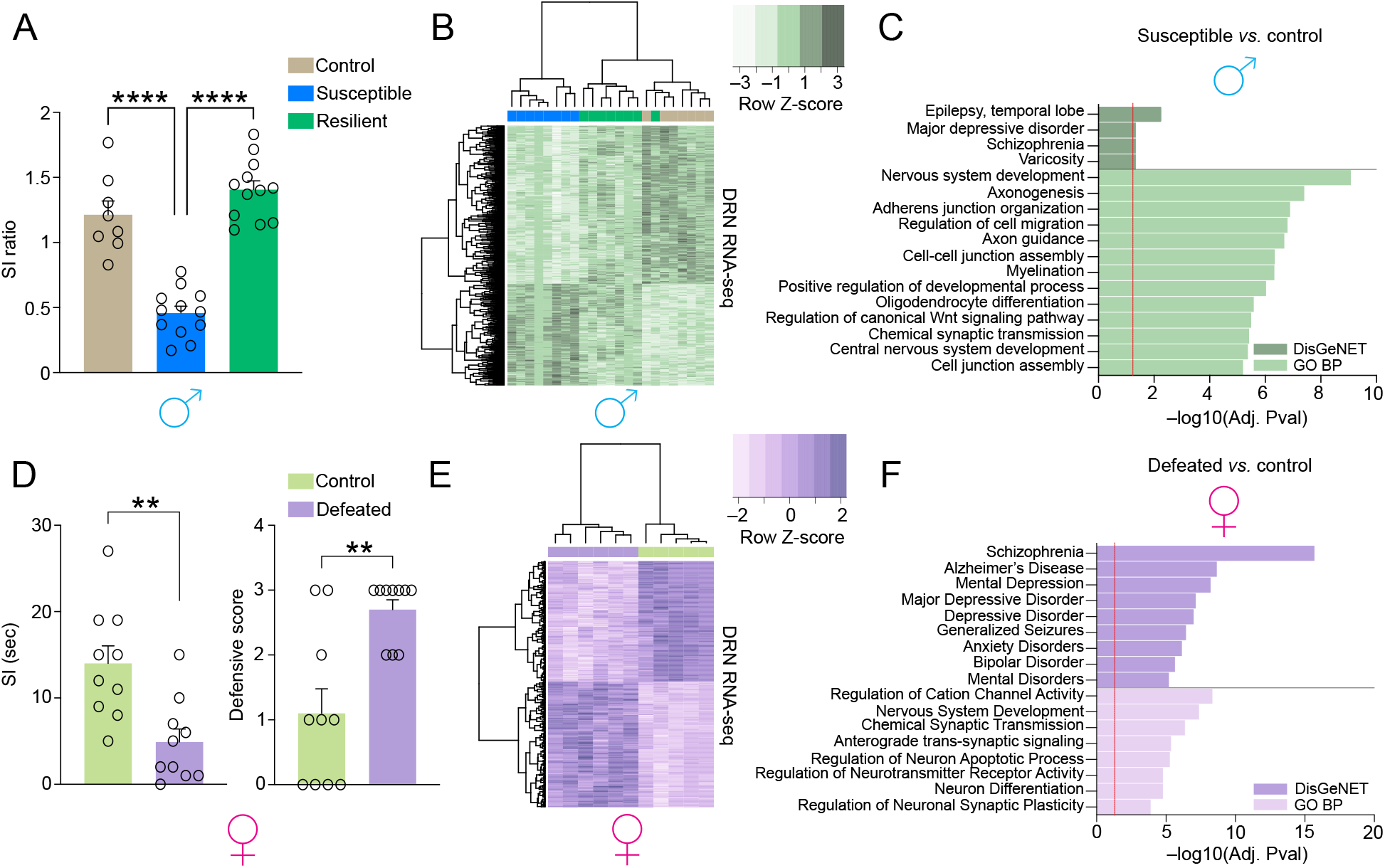
Chronic social stress in both males and female mice results in altered gene expression programs in DRN. **(A)** Social interaction ratio of control *vs.* stress-susceptible *vs.* resilient male mice (*n* = 8 in controls, 12 in susceptible and resilient groups). Data were analyzed using a one-way ANOVA, with significant main effects (*p*<0.0001, F_2,29_ = 53.44). Tukey’s multiple comparisons test revealed significant differences between control *vs.* susceptible mice (*p*<0.0001) and susceptible *vs.* resilient mice (*p*<0.0001). **(B)** Clustering of control, susceptible and resilient groups for 1,502 differentially expressed (DE) genes (susceptible *vs.* control; *n* = 7-8/group, FDR<0.05). **(C)** Example GO Biological Process and DisGeNET pathway enrichment (FDR<0.05) for the PCGs differentially expressed (at FDR<0.1) in susceptible *vs.* control males. Dashed line indicates significance via adjusted p-value. **(D_Left_)** Social interaction time of control *vs.* socially defeated female mice (*n* = 10/group). Data were analyzed via Student’s two-tailed *t*-test with significant differences observed between defeated *vs.* control mice (*p*=0.0021, t_18_ = 3.582). **(D_Right_)** Defensive score of control *vs.* socially defeated female mice. Data were analyzed via Mann-Whitney test (unpaired) with significant differences observed between defeated *vs.* control mice (*p*=0.0034, *U*=14.50). **(E)** Clustering of defeat and control groups for 234 DE genes (defeat *vs.* control; *n* = 5-6/group, FDR< 0.05). **(F)** Example GO Biological Process and DisGeNET pathway enrichment (FDR<0.05) for the PCGs differentially expressed in defeat *vs.* control females (at FDR< 0.1). Dashed line indicates significance via FDR. For all bar graphs, data presented as mean ± SEM.

Given the vast literature indicating prominent sex differences with respect to disparities of onset, lifetime prevalence and symptomatic presentation of MDD in humans (57–59), as well as stress vulnerability phenotypes in preclinical animal models (60), we next sought to examine gene expression programs in DRN of chronically stressed (i.e., defeated) female mice in order to compare to those transcriptional patterns observed in males. To do so, we performed a recently established CSDS paradigm in female mice that similarly recapitulates numerous features of MDD, as well as behaviors observed in male rodents following CSDS (41,61), including increased social avoidance and heightened levels of defensive behaviors (**Fig. 1D**). In our paradigm, defeated females were found to be entirely susceptible to CSDS, so we therefore compared defeated females to susceptible males in subsequent analyses. Following CSDS, we again performed bulk RNA-seq on DRN tissues from control *vs.* defeated female mice, followed by differential expression analysis to compare the two groups. We found that defeated females exhibited significant alterations in the expression of 339 PCGs (FDR<0.1) *vs.* respective controls – far fewer than that observed in males following CSDS – with unsupervised clustering revealing a clear pattern of separation between subjects by ‘treatment’ type (**Fig. 1E, Supplemental Data Table 10**). While a more modest number of loci were found to be dysregulated in female *vs.* male DRN, gene set enrichment analyses (GO Biological Process and DisGeNET) again identified significant (FDR<0.05) overlaps with shared pathways/processes involved in nervous system development, chemical synaptic transmission and psychiatric/mood-related disorders (e.g., MDD depressive disorder, anxiety disorders, bipolar disorder, etc.) (**Fig. 1F, Supplemental Data Tables 11-12**). Again, downregulated genes in females were found to contribute most significantly to gene ontologies observed when assessing the dataset irrespective of directionality (FDR<0.05; **Fig. SC-D, Supplemental Data Tables 13-16**). In addition, while male susceptible mice clearly displayed more robust alterations in gene expression *vs.* defeated females, a subset of these dysregulated loci were found to significantly overlap between the sexes (odds ratio = 3.1; p = 1.3e-16) (**Fig. S1E**), with these shared genes also displaying significant enrichment for pathways/processes involved in brain development, synaptic transmission and mood-related disorders (FDR<0.05; **Fig. S1F, Supplemental Data Tables 17-18**). These data indicate that, as in stress-susceptible males, gene expression programs in defeated female DRN also appear relevant to abnormal neuronal and behavioral plasticity associated with affective disturbances.

### H3 serotonylation is altered genome-wide in DRN of male and female mice acutely following chronic social stress

Given the transcriptional responsiveness of DRN to chronic stress exposures in both male and female mice, we next sought to interrogate potential chromatin-related mechanisms that may contribute to these observed dynamics. Since DRN is enriched for 5-HT-producing neurons, a monoaminergic cell population that also displays robust enrichment for histone H3 serotonylation (18), we further explored potential regulation of H3K4me3Q5ser dynamics in DRN of mice 24 hr following social interaction (SI) testing. Using western blotting to first assess global levels of the combinatorial PTM, we found that H3K4me3Q5ser was nominally (p≤0.05, unpaired t test) downregulated in male DRN when comparing stress-susceptible *vs.* control animals, an effect that was not observed in stress-resilient *vs.* control comparisons (**Fig. 2A**). However, when comparing all three groups (one-way ANOVA), we observed robust differences between stress-susceptible *vs.* stress-resilient animals, with stress-susceptible mice displaying significant deficits in the mark. Similarly, when comparing female defeated *vs.* control animals 24 hr after CSDS, we found that the serotonylation mark was also downregulated (**Fig. 2B**). To assess whether these changes may be clinically relevant, we next measured H3K4me3Q5ser levels in postmortem DRN of humans with MDD *vs.* demographically matched controls, where we found that the serotonylation mark was also downregulated in MDD individuals without ADs present at their time of death (**Fig. 2C, left;** p<0.05, unpaired t test). Interestingly, however, when comparing levels of the mark in DRN from humans diagnosed with MDD with ADs present at their time of death, we observed that H3K4me3Q5ser levels were similar to those of their respectively matched controls (**Fig. 2C, right;** p>0.05, unpaired t test), suggesting an interaction between the mark’s expression, MDD diagnosis and AD exposures.

**Fig. 2.**
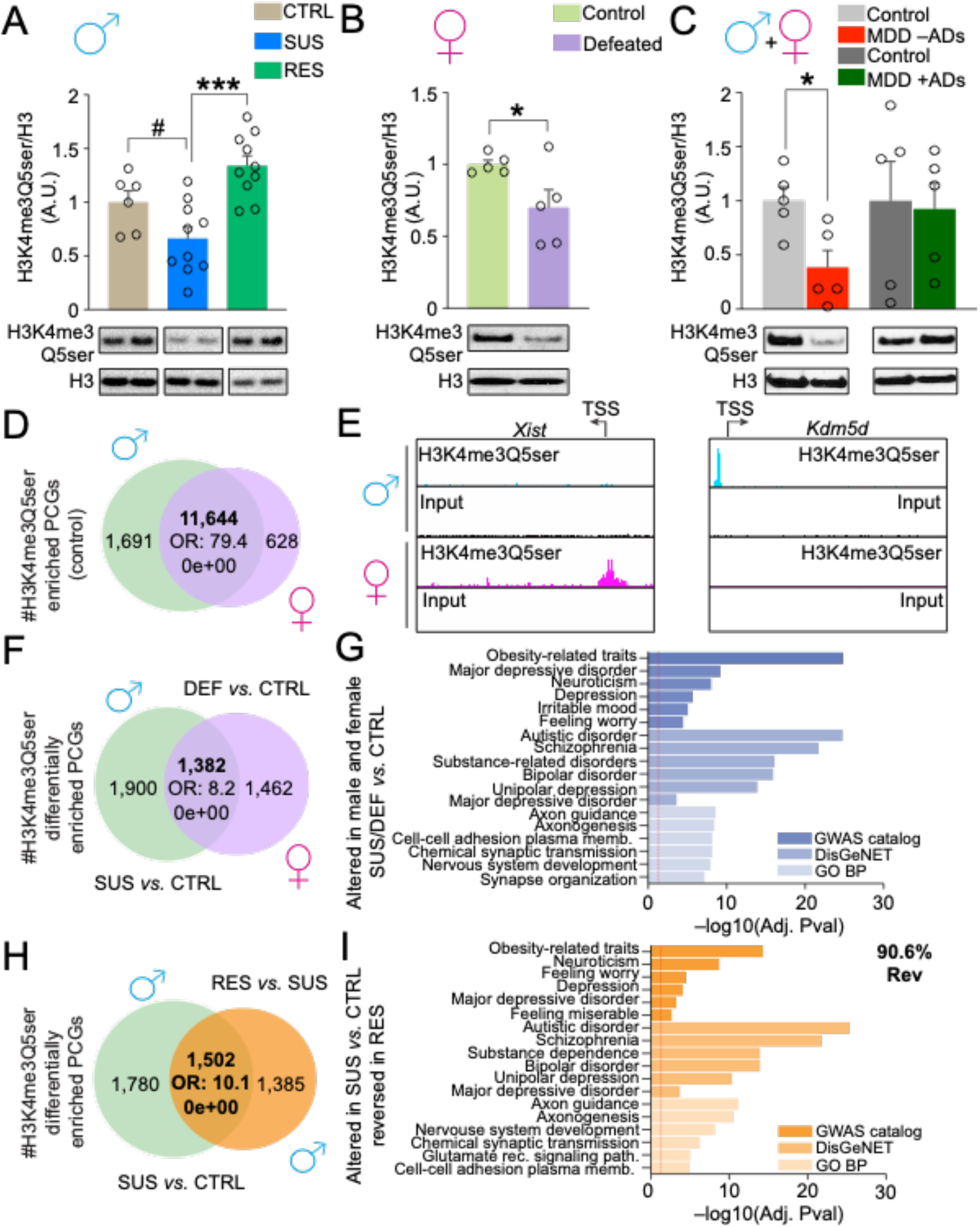
Chronic social stress promotes altered H3 serotonylation dynamics in DRN. **(A)** Western blotting analysis of H3K4me3Q5ser in DRN (24hr post-SI testing) of control *vs.* stress-susceptible *vs.* resilient male mice (*n* = 6-10/group). Data were analyzed using a one-way ANOVA, with significant main effects observed (*p*=0.0002, F_2,23_ = 12.43). Tukey’s multiple comparisons test revealed significant differences between susceptible *vs.* resilient mice (*p*=0.0001), and an *a posteriori* Student’s two-tailed *t*-test revealed a significant difference between control *vs.* susceptible mice (*p*=0.05, t_14_ = 2.117). Total histone H3 levels were used as a loading control. **(B)** Analysis of H3K4me3Q5ser in DRN of control *vs.* socially defeated female mice (*n* = 5/group). Data were analyzed via a Student’s two-tailed *t*-test with a significant difference observed between defeated *vs.* control mice (*p*=0.0473, t_8_ = 2.341). Total histone H3 levels were used as a loading control. **(C)** Analysis of H3K4me3Q5ser in DRN from human postmortem brain of MDD individuals with or without antidepressants onboard at their time of death *vs.* respective demographically matched controls (*n* = 5/group). Data were analyzed via Student’s two-tailed *t*-tests (individual MDD groups *vs.* matched controls) with significant differences observed between MDD –AD’s *vs.* age/sex matched controls (*p*=0.0166, t_8_ = 3.020). Total histone H3 levels were used as a loading control. For all western blotting graphs, */#*p* <0.05, \*\*\**p* <0.001. Data presented as mean ± SEM. A.U., arbitrary units, normalized to respective controls. **(D)** Venn diagram depicting the degree of overlap between H3K4me3Q5ser enriched PCGs (peaks >5 fold-enriched over input, FDR<0.05) in control male *vs.* female DRN (*n* = 3/group, 3-4 samples pooled per *n*). Odds ratio (OR) and respective p-value of overlap is provided. **(E)** Representative IGV genome-browser tracks for two sex-specific loci (*Xist* in female DRN and *Kdm5d* in male DRN) displaying sex specific enrichment of permissive H3K4me3Q5ser *vs.* respective inputs. **(F)** Venn diagram depicting the degree of overlap between male *vs.* female PCGs displaying differential enrichment for H3K4me3Q5ser in DRN as a consequence of CSDS (male susceptible *vs.* control, and female defeated *vs.* control; *n* = 3/group, 3-4 samples pooled per *n*, FDR<0.05, Log2FC ≥1.5 or ≤–1.5). Odds ratio (OR) and respective p-value of overlap is provided. **(G)** Selected GWAS catalog, DisGeNET and GO Biological Process pathways for PCGs displaying overlapping (male *vs.* female; 1,382 PCGs) differential enrichment for H3K4me3Q5ser as a consequence of CSDS (FDR<0.05). **(H)** Venn diagram depicting the degree of overlap between male stress-susceptible *vs.* control and male stress-resilient *vs.* stress-susceptible PCGs displaying altered H3K4me3Q5ser enrichment in DRN (*n* = 3/group, 3-4 samples pooled per *n*), FDR<0.05, Log2FC ≥1.5 or ≤–1.5). Odds ratio (OR) and respective p-value of overlap is provided. **(I)** Selected GWAS catalog, DisGeNET and GO Biological Process pathways for PCGs displaying overlapping and reversed (90.6% of overlapping PCGs) differential enrichment for H3K4me3Q5ser in male stress-susceptible *vs.* control and male stress-resilient *vs.* stress-susceptible comparisons (FDR<0.05). See **Fig. S7A-C** for uncropped blots.

Next, to assess whether alterations in global levels of H3K4me3Q5ser correspond with meaningful patterns of genomic regulation following chronic stress exposures, we performed ChIP-seq for the mark in DRN of both males (control *vs.* susceptible *vs.* resilient) and females (control *vs.* defeated) 24 hr after SI testing. Following peak calling (FDR<0.05, >5 fold-enrichment over input, **Supplemental Data Tables 19-23**), we first assessed the degree of overlap between PCGs enriched for H3K4me3Q5ser in control males *vs.* females, where we observed a significant degree of overlap between the sexes (odds ratio = 79.4; p = 0e+00); **Fig. 2D**), with non-overlapping peaks largely representing sex-specific loci (e.g., *Xist* in females and *Kdm5d* in males; **Fig. 2E**). Note that the majority of peaks identified in both male and female DRN were found to be located throughout genic loci, particularly within gene promoters and often enriching proximally to transcriptional start sites (**Fig. S2A-B**). We then performed differential enrichment analysis for the mark (FDR<0.05, Log2FC ≥ 1.5 or ≤ –1.5, **Supplemental Data Tables 24, 27**) at peak-enriched PCGs (the most common sites of differential enrichment in both males and females (**Fig. S2C-D**) comparing the degree of overlap between differentially enriched loci between stress-susceptible males *vs.* defeated females (*vs.* their respective controls), where we observed that ∼42.1% and ∼48.6% of differentially enriched PCGs overlapped (odds ratio = 8.2; p = 0e+00) between males and females, respectively, following chronic stress (**Fig. 2F**). Similar to the gene expression results presented in **Figure 1**, these overlapping gene sets between male and female stress-exposed mice were demonstrated to enrich (FDR<0.05) for pathways/biological processes (GO Biological Process) associated with neuronal development and synaptic regulation, as well as disease-enriched (GWAS catalog, DisGeNET) loci related to psychiatric, neurodevelopmental and affect-related disorders (e.g., MDD, Irritable Mood, Feeling Worry, Bipolar Depression, Unipolar Depression, etc.) (**Fig. 2G, Supplemental Data Tables 28-30**). Importantly, a subset of these differentially enriched loci were observed to significantly overlap with genes demonstrated to be differentially expressed in response to stress in both males (odds ratio = 2.0; p = 1.1e-27) and females (odds ratio = 1.8; p = 4.5e-07) (**Fig. S2E-F**), with these overlapping PCGs also displaying significant enrichment (FDR<0.05) for disease pathways (DisGeNET) associated with MDD, mood disorders, bipolar disorder, unipolar depression, etc. in both sexes (**Fig. S2G**, **Supplemental Data Tables 34-35**). Finally, given our earlier western blotting results in males showing that decreased H3K4me3Q5ser levels in susceptible animals were not observed in resilient mice, we next examined the degree of overlap (odds ratio = 10.1; p = 0e+00) between differentially enriched PCGs in stress-susceptible *vs.* stress-resilient mice, finding that ∼90.6% of PCGs exhibiting dynamics in susceptible mice displayed reversal in these enrichment patterns in stress-resilient animals (**Fig. 2H, Supplemental Data Tables 25-26**). Subsequent gene set enrichment analyses (FDR<0.5) again revealed strong associations between those loci displaying reversals in enrichment between susceptible *vs.* resilient mice and pathways/processes (GO Biological process) related to neurodevelopmental processes and synaptic organization/function, along with significant enrichment in disease associated pathways (GWAS catalog, DisGeNET) related to affective disorders (e.g., Depression, Feeling Worry, MDD, Bipolar Depression, etc.) and other psychiatric syndromes (**Fig. 2I, Supplemental Data Tables 31-33**). In sum, our genomic data acutely following CSDS indicated that alterations in H3K4me3Q5ser enrichment patterns in DRN in response to chronic stress significantly correlate with abnormal transcriptional programs associated with MDD and other affective disorders.

### Chronic AD treatments reverse stress susceptibility and rescue stress-induced H3 serotonylation dynamics in DRN

Considering that our western blotting data in human DRN revealed that global levels of H3K4me3Q5ser were altered in individuals with MDD without ADs onboard at their time of death, an effect that was not observed in patients with ADs onboard at their time of death *vs.* respectively matched controls, we next sought to explore whether the mark may similarly be responsive to chronic AD treatments following CSDS in mice. To examine this, male mice were subjected to 10 days of CSDS, assessed for SI and separated into control *vs.* susceptible *vs.* resilient populations (**Fig. 3A-C**; Pre-treatment) before being treated for 30 days with the SSRI AD fluoxetine *vs.* water as a vehicle control (42) (**Fig. 3A-C**; Post-treatment). Following another round of SI testing to examine behavioral reversal of the susceptibility phenotype in previously susceptible mice, DRN tissues were collected for western blotting analysis of H3K4me3Q5ser. As expected, susceptible mice remained susceptible, as measured via SI, following chronic treatments with water (**Fig. 3B**). However, susceptible animals treated with chronic fluoxetine displayed significant reversal of previously observed SI deficits (**Fig. 3C**). Using this protracted timeline, which may better reflect the persistence of stress-vulnerable states *vs.* examinations 24 hr post-CSDS (as in **Figure 2**), we no longer observed a global downregulation of H3K4me3Q5ser – a phenomenon that was seen acutely following chronic stress – but rather found that the mark accumulates in DRN of stress-susceptible mice treated with water *vs.* vehicle treated control and stress-resilient animals. (**Fig. 3D**). This accumulation, however, was found to be significantly attenuated by chronic fluoxetine treatments in stress-susceptible mice, with levels of the mark normalizing to those of both control and resilient animals; fluoxetine administration did not impact levels of the mark in control or resilient mice, animals that remained behaviorally unaffected in response to chronic AD treatments. These data demonstrated that behavioral responsiveness to ADs following chronic stress in susceptible mice (but not in the absence of stress or in resilient animals) corresponds with reductions in H3K4me3Q5ser levels in DRN, perhaps suggesting a role for AD-mediated H3K4me3Q5ser downregulation in the alleviation of stress-induced behavioral deficits.

**Fig. 3.**
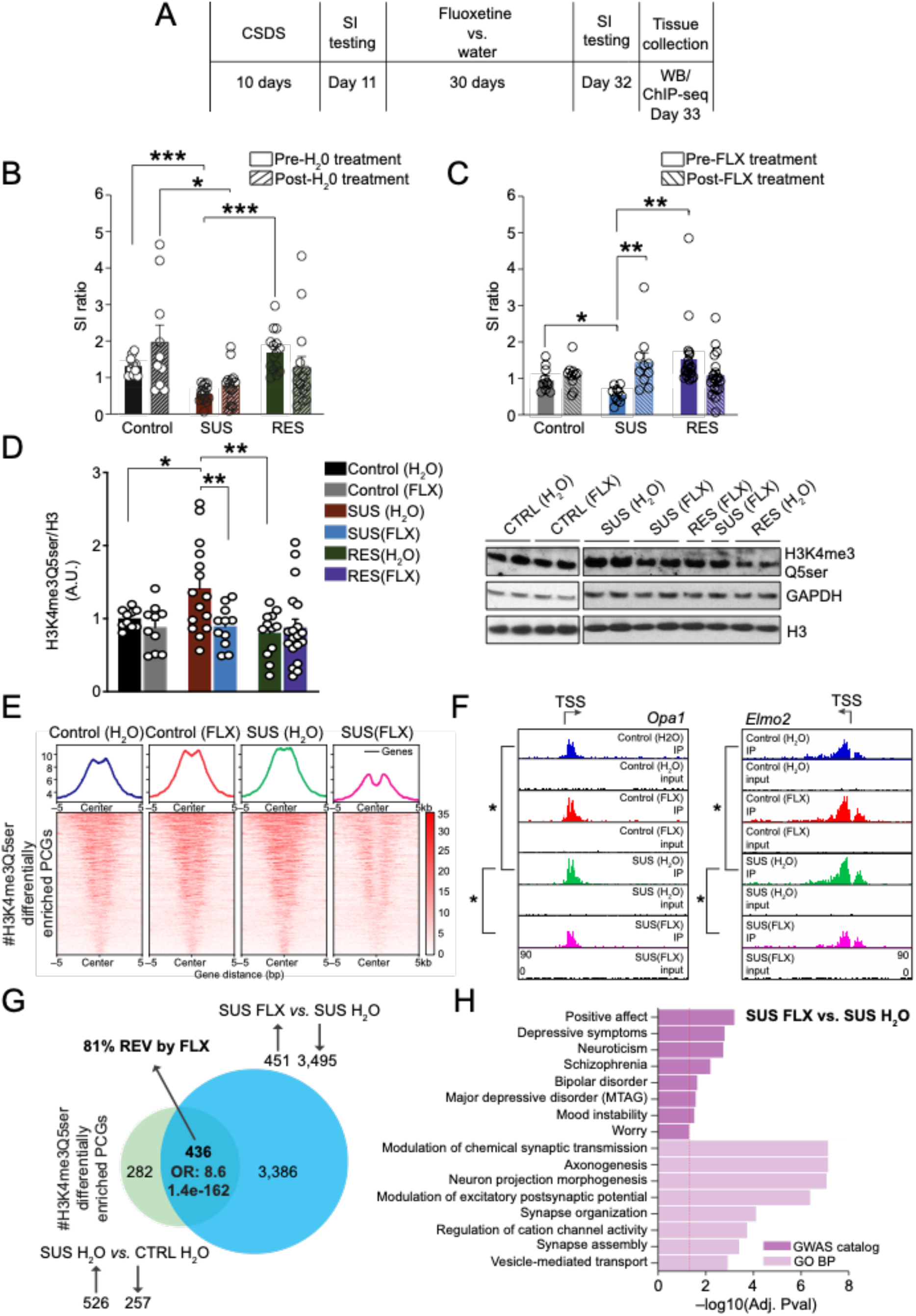
Chronic fluoxetine treatments rescue behavioral deficits and stress-induced H3K4me3Q5ser dynamics in DRN in stress-susceptible male mice. **(A)** Schematic of experimental timeline for CSDS and SI testing in male mice, followed by 30 days of fluoxetine treatments *vs.* water, additional SI testing and DRN collections. **(B)** SI ratio of control *vs.* stress-susceptible *vs.* stress-resilient male mice, pre-*vs.* post-30 days of water administration as a vehicle control (*n* = 10-15/group). Data were analyzed using a two-way repeated measures ANOVA, with significant main effects of stress (*p*=0.0005, F_2,37_ = 9.298) and interaction of stress x time (*p*=0.0234, F_2,37_ = 4.162) observed. Posthoc *t*-tests with Bonferroni correction revealed significant differences between control versus stress-susceptible mice, pre-treatment (*p*=0.0003), stress-susceptible versus resilient mice, pre-treatment (*p*=0.0003), and control versus stress-susceptible mice, post-treatment (*p*=0.0201 **(C)** SI ratio of control *vs.* stress-susceptible *vs.* resilient male mice, pre-*vs.* post-30 days of fluoxetine administration (*n* = 10-19/group). Data were analyzed using a two-way repeated measures ANOVA, with significant main effect of interaction of stress x treatment (*p*=0.0018, F_2,36_ = 7.548). Bonferroni’s multiple comparisons tests revealed significant difference between stress-susceptible mice, pre-*vs.* post-30 days of fluoxetine administration (*p*=0.0098). Neither control (*p* > 0.9999) nor resilient mice (*p*=0.0917) displayed significant alteration in SI following fluoxetine treatments. Posthoc *t*-tests with Bonferroni correction revealed significant differences between control versus stress-susceptible mice, pre-treatment (*p*=0.0111) and stress-susceptible versus resilient mice, pre-treatment (*p*=0.0066). **(D)** Analysis of H3K4me3Q5ser/H3 in DRN of control *vs.* stress-susceptible *vs.* stress-resilient male mice following 30 days of fluoxetine administration *vs.* water (*n* = 10-19/group). Data were analyzed using a two-way ANOVA, with significant main effects of stress (*p*=0.0289, F_2,71_ = 3.725) and stress x fluoxetine (*p*=0.0420, F_2,71_ = 3.316). Sidak’s multiple comparisons tests revealed significant differences between stress-susceptible mice post-30 days of fluoxetine administration *vs.* stress-susceptible mice post-30 days of water administration (*p*=0.0094), and Tukey’s multiple comparisons tests revealed significant differences between stress-susceptible mice post-30 days of water administration *vs.* control mice post-30 days of water administration (*p*=0.0554), and stress-susceptible mice post-30 days of water administration *vs.* resilient mice post-30 days of water administration (*p*=0.0013). To control for samples spread across numerous membranes, GAPDH levels were used as an additional loading control. **(E)** Peak-centered heatmaps depicting H3K4me3Q5ser enrichment at PCGs displaying significant differential enrichment (FDR<0.05, Log2FC ≥1.0 or ≤–1.0) between SUS FLX *vs.* SUS H_2_O mice for each of the four groups (control H_2_O, control FLX, SUS H_2_O and SUS FLX). **(F)** Representative IGV genome-browser tracks for two genic loci (*Opa1* and *Elmo2*) displaying significantly (*) increased enrichment for H3K4me3Q5ser in SUS (H_2_O) *vs.* control (H_2_O) mice and rescue following of this increased enrichment in SUS (FLX) *vs.* SUS (H_2_O) animals (respective inputs are included). **(G)** Venn diagram depicting the degree of overlap between male PCGs displaying protracted differential enrichment of H3K4me3Q5ser in DRN as a consequence of CSDS (susceptible H_2_O *vs.* control H_2_O) *vs.* PCGs displaying regulation of the mark by chronic fluoxetine treatments in susceptible mice (susceptible FLX *vs.* susceptible H_2_O); *n* = 3/group, 3-4 samples pooled per *n*, FDR<0.05, Log2FC ≥1.0 or ≤–1.0). Odds ratio (OR) and respective p-value of overlap is provided. **(H)** Selected GWAS catalog and GO Biological Process pathways for PCGs displaying differential enrichment for H3K4me3Q5ser in male stress-susceptible FLX *vs.* stress-susceptible H_2_O comparisons (FDR<0.05). For all bar graphs, \**p* < 0.05, \*\**p* < 0.01, \*\*\**p* < 0.001. Data presented as mean +/-SEM. A.U., arbitrary units, normalized to respective controls. See **Fig. S7D** for uncropped blots.

Given these western blotting results, we next aimed to explore H3K4me3Q5ser dynamics genome-wide at this protracted timepoint following CSDS −/+ chronic fluoxetine treatments. Following ChIP-seq for the mark in DRN tissues from male control *vs.* stress-susceptible mice – vehicle (H_2_O) *vs.* fluoxetine – we first assessed the degree of overlap between differentially enriched PCGs regulated by chronic stress at acute (stress-susceptible *vs.* control, 24 hr post-CSDS) *vs.* protracted (stress-susceptible H_2_O *vs.* control H_2_O, 30 d post-CSDS) periods following CSDS. While we found that a greater amount of genes displayed stress-induced H3K4me3Q5ser dynamics (FDR<0.05, Log2FC ≥ 1.0 or ≤ –1.0; note that a slightly lower Log2FC cutoff was used in these comparisons *vs.* those in **Fig. 2** to account for batch variability between experiments) at acute (3,879) *vs.* protracted (718) periods following CSDS in the absence of fluoxetine (the majority of which for both comparisons displayed increased enrichment of the mark), 223 of these PCGs were found to significantly overlap between the two timepoints (odds ratio = 2.3; p = 1.4e-21); **Fig. S3A, Supplemental Data Tables 36-39**), with many of these overlapping genes (75%) displaying consistent patterns of regulation by CSDS (127 up/up, 41 down/down). Furthermore, gene set enrichment analyses (FDR<0.5) of these 223 overlapping PCGs revealed strong associations with pathways/processes (GO Biological process) related to synaptic organization/function, along with significant enrichment in disease associated pathways (GWAS catalog) related to affective disorders (e.g., MDD, depressive symptoms) and other psychiatric syndromes (**Fig. S3B, Supplemental Data Tables 40-41**). We next aimed to assess the impact of chronic fluoxetine exposures, which effectively reversed stress-susceptibility and stress-induced gene expression (see **Fig. 3B-C, Fig. S3C, Supplemental Data Tables 42-43**), on H3K4me3Q5ser dynamics genome-wide in mouse DRN. In doing so, we found that 81% of overlapping PCGs (comparing susceptible H_2_O *vs.* control H_2_O and susceptible fluoxetine *vs.* susceptible H_2_O gene lists) displaying differential enrichment of the mark at protracted periods following CSDS exhibited restoration of these dynamics in response to chronic fluoxetine treatments (odds ratio = 8.6; p = 1.4e-162), with fluoxetine exposures additionally resulting in a robust loss of H3K4me3Q5ser enrichment at a large number of genes (3,495) that were not observed to be regulated in their serotonylation state by chronic stress alone (i.e., an apparent interaction between stress x fluoxetine was clearly observed; **Fig. 3E-G, Supplemental Data Table 44-46**); note that many fewer PCGs were found to display significant H3K4me3Q5ser dynamics in DRN as a consequence of chronic fluoxetine exposures in control mice, which is consistent with a lack of behavioral responsiveness to ADs in these animals (**Supplemental Data Table 47**). Again, gene set enrichment analyses (FDR<0.5) of PCGs displaying altered enrichment in stress-susceptible fluoxetine *vs.* stress-susceptible H_2_O mice revealed strong associations with pathways/processes (GO Biological process) related to synaptic organization/function, along with significant enrichment in disease associated pathways (GWAS catalog) related to mood disorders (e.g., MDD, bipolar disorder, mood instability) and other psychiatric syndromes (**Fig. 3H, Supplemental Data Tables 48-49**). These data suggested that while fluoxetine indeed functions, at least in part, to reverse stress-induced H3K4me3Q5ser dynamics in DRN, it also serves to promote more global alterations (predominantly reduced enrichment) of the mark, which may additionally contribute to reversals in stress-susceptibility. Next, given that >90% of overlapping stress-regulated genes between stress-resilient *vs.* stress-susceptible mice 24 hr post-CSDS were found to display opposing patterns of H3K4me3Q5ser regulation (**Fig. 2H**), we next sought to assess whether PCGs displaying altered H3K4me3Q5ser enrichment in stress-susceptible mice + chronic fluoxetine may overlap with PCGs displaying reversals in the mark’s enrichment in stress-resilient *vs.* stress-susceptible comparisons (FDR<0.05, Log2FC ≥ 1.0 or ≤ –1.0, **Supplemental Data Table 50**). Indeed, we identified significant overlaps between these two comparisons (**Fig. S3D**, odds ratio = 2.5; p = 4.8e-90), with the overlapping genes significantly enriching (FDR<0.05) for pathways/processes (GO Biological process) related to synaptic organization/function, along with significant enrichment in disease associated pathways (GWAS catalog) related to mood disorders (e.g., MDD, bipolar disorder, positive affect) and other psychiatric illnesses (**Fig. S3E, Supplemental Data Tables 51-52**).

Finally, to examine whether fluoxetine-induced changes in H3K4me3Q5ser enrichment that were observed in stress-susceptible mice correlate with genes displaying similar patterns of regulation in human brain, we next performed ChIP-seq for the mark in human postmortem DRN tissues from individuals diagnosed with MDD −/+ ADs onboard at their time of death (FDR<0.05, Log2FC ≥ 1.0 or ≤ –1.0, **Supplemental Data Tables 53-58**). While only a handful of differentially enriched PCGs were observed when comparing MDD – ADs *vs.* matched controls – likely owing to the heterogenous nature of MDD and our limited sample size – such assessments did identify significant and ontologically relevant (FDR<0.05, **Supplemental Data Tables 59-60**) overlaps between PCGs displaying altered dynamics of the mark in susceptible fluoxetine *vs.* susceptible H_2_O mice and in human subjects with MDD + *vs.* – ADs (**Fig. S3F-G**, odds ratio = 2.2; p = 4.4e-92); note that a greater number of PCGs in total displayed loss (4,146) *vs.* gain (3,034) of the mark in MDD patients + *vs. –* ADs, data which are consistent with our fluoxetine findings in mice. These data suggest that alterations in H3K4me3Q5ser enrichment observed in behaviorally responsive, fluoxetine treated CSDS mice may be clinically relevant and may reflect functionally important chromatin adaptations that occur in human MDD subjects undergoing AD treatments.

### Directly reducing H3 serotonylation in DRN promotes stress resilience through reversal of stress-mediated gene expression programs

Since we observed that H3K4me3Q5ser levels were elevated in DRN of susceptible *vs.* resilient mice over protracted periods following chronic stress exposures, a phenomenon that was largely reversed by ADs, we next aimed to explore whether prophylactically reducing H3 serotonylation in DRN may prevent the precipitation of stress-mediated gene expression programs and/or behavioral susceptibility. To examine this, male mice were injected intra-DRN with one of three lentiviral vectors – which transduce both neurons and glia, as all cell-types in DRN have previously been shown to express the serotonylation mark (18); thus, we aimed to express these constructs in a non-cell-type restrictive manner – expressing either GFP (aka empty) or H3.3 WT controls *vs.* H3.3Q5A, the latter of which functions as a dominant negative by incorporating into neuronal chromatin without being able to be monoaminylated, thereby reducing levels of H3 serotonylation at affected loci (**Fig. S4A-D**) (18). Following viral transduction and recovery, mice underwent CSDS and then were assessed via SI testing to examine avoidance behavior, after which time, virally transduced DRN tissues were collected for RNA-seq analysis (**Fig 4A**). A separate cohort of mice were surgerized to validate the efficiency of H3.3 incorporation into chromatin in DRN via immunohistochemistry/immunofluorescence (**Fig. 4B**). Following CSDS in virally transduced animals, we observed significant deficits in social interaction in both viral control groups (empty and H3.3 WT, neither of which impact H3 serotonylation – see **Fig. S4**; note that due to the experimental design of this experiment, susceptible and resilient behavioral readouts occurred post viral manipulations). However, we found that reducing H3 serotonylation in DRN using the dominant negative H3.3Q5A virus rescued CSDS-induced social avoidance behavior (in effect increasing the proportion of resilient animals observed post-CSDS *vs.* empty or H3.3 WT groups), indicating that viral-mediated downregulation of H3K4me3Q5ser in chronically stressed animals is sufficient to promote behavioral resilience (**Fig. 4C**). And while our AD data presented in **Figure 3** could not definitively link observed fluoxetine-induced reductions in H3K4me3Q5ser to the reversals of stress susceptibility observed post-AD treatment, those findings were indeed consistent with our viral manipulation experiments, which causally linked inhibition of the mark during stress exposures to the promotion of stress-resilience. Expression of H3.3Q5A in DRN did not affect SI behavior in control (i.e., non-CSDS) mice; however, attenuation of H3 serotonylation in non-stressed mice was found to decrease behavioral despair in the forced swim test (FST) (**Fig. S5A**), with no impact of viral manipulations observed in anxiety-related tasks, such as the elevated plus maze (EPM; **Fig. S5B**) or open field test (OFT; **Fig. S5C**).

**Fig. 4.**
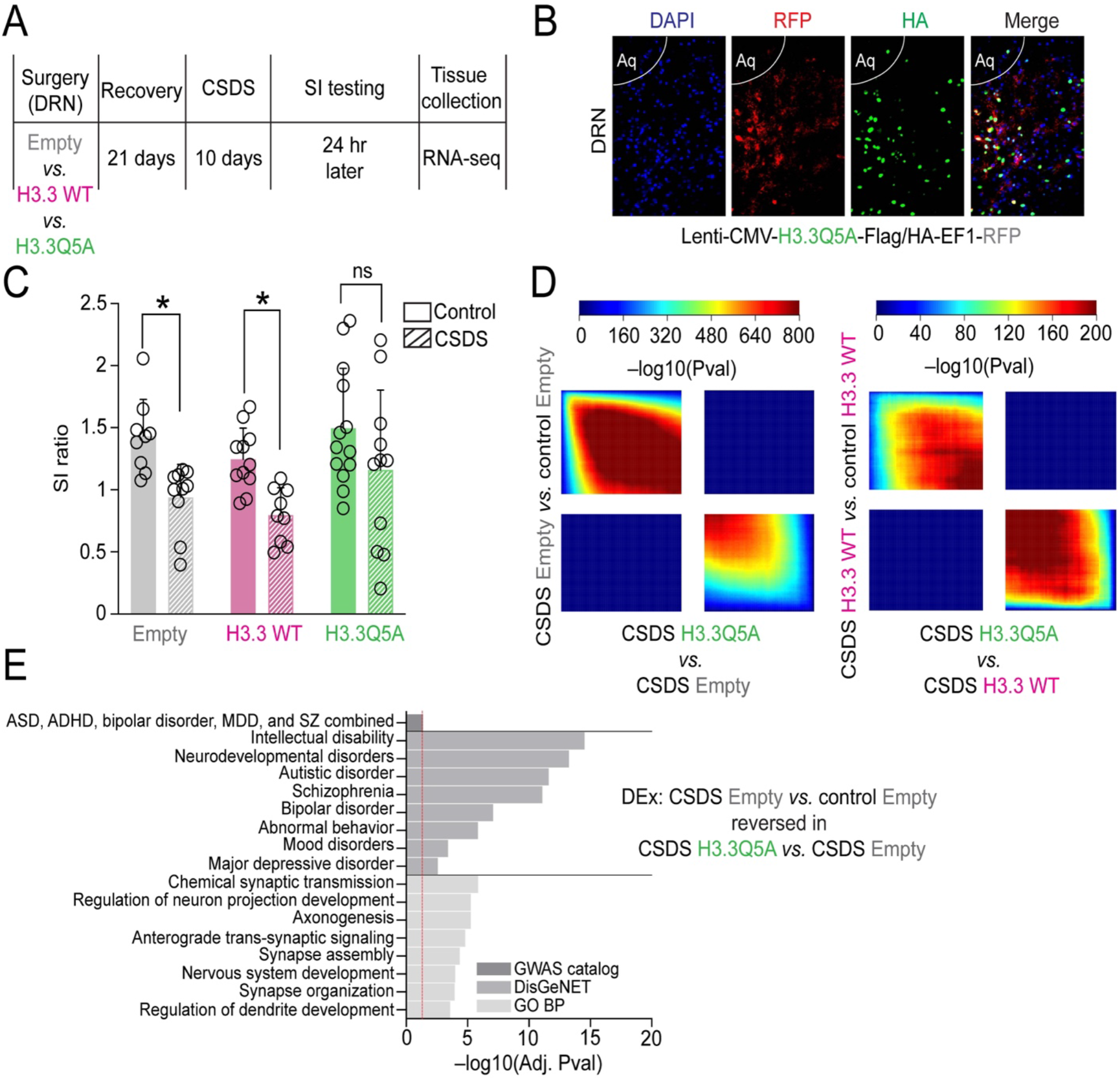
Viral-mediated downregulation of H3 serotonylation in DRN promotes stress resilience and rescues stress-induced gene expression. **(A)** Schematic of experimental timeline for male CSDS after intra-DRN viral transduction by empty vector, H3.3 WT or H3.3Q5A vectors, followed by behavioral testing and tissue collections for RNA-seq. **(B)** Representative IHC/IF images of mouse DRN transduced with a lentivirus expressing H3.3Q5A-HA-EF1-RFP (red fluorescent protein) overlayed with staining for HA and a nuclear co-stain [4′,6-diamidino-2-phenylindole (DAPI)]. **(C)** SI ratios of GFP, H3.3 WT and H3.3Q5A transduced mice, control *vs.* CSDS (*n* = 9-13/group). Data were analyzed using a two-way ANOVA, with significant main effects of stress observed (*p*=0.0001, F_1,57_ = 17.29). Bonferroni’s multiple comparisons tests revealed significant differences between control *vs.* CSDS groups in GFP (*p*=0.0310) and H3.3 WT mice (*p*=0.0474), with no differences observed between control *vs.* CSDS H3.3Q5A mice. **(D)** Threshold-free RRHO analyses comparing transcriptional profiles for stress-regulated genes in empty vector and H3.3 WT-transduced DRN (control *vs.* CSDS) to H3.3Q5A-transduced DRN from CSDS mice (*n* = 4-9/group), demonstrating that H3.3Q5A significantly reversed gene expression programs observed in response to stress in both control groups. Each pixel represents the overlap between differential transcriptomes, with the significance of overlap of a hypergeometric test color-coded. **(E)** Selected GWAS catalog, DisGeNET and GO Biological Process pathways for PCGs displaying differentially expressed genes in CSDS empty *vs.* control empty comparisons and rescue in CSDS H3.3Q5A *vs.* CSDS Empty comparisons (FDR<0.1). Select enriched pathways are shown (FDR<0.05).

Next, to examine whether behavioral rescue in H3.3Q5A-expressing mice may correspond to a restoration of gene expression abnormalities elicited by chronic stress exposures, we performed bulk RNA-seq on microdissected, virally transduced DRN tissues from control *vs.* CSDS mice. Rank-rank hypergeometric overlap (RRHO) analysis revealed, in comparison to gene expression programs potentiated by chronic stress in both control groups (empty – left, H3.3 WT – right), that transduction by H3.3Q5A significantly reversed stress-induced gene expression profiles (**Fig. 4D, Supplemental Data Tables 61-64**). Importantly, gene expression programs found to be induced by CSDS in virally transduced animals (e.g., empty vector) significantly correlated with differential gene expression patterns observed in susceptible *vs.* control comparisons (24 hr post-SI testing) using tissues from non-virally transduced mice (from **Figure 2**), with H3.3Q5A manipulations similarly reversing the expression of these stress impacted genes (**Fig. S6A-B**). These data demonstrate that H3 serotonylation is important for potentiating stress-associated patterns of transcriptional dysregulation in DRN, abnormalities that may contribute importantly to the behavioral deficits observed. Finally, to elucidate the specific gene sets and biological pathways that may be affected by H3K4me3Q5ser downregulation in stress-susceptible animals, we performed differential expression analysis comparing H3.3Q5A *vs.* empty-expressing mice −/+ CSDS, and then used the list of significantly rescued genes following H3.3Q5A manipulations (**Fig. S6C-D**) to perform gene ontology analyses (**Fig. 4E, Supplemental Data Tables 65-67**). These genes were subjected to gene set enrichment analysis (GWAS catalog, DisGeNET and GO Biological process), which significantly implicated phenotypic and disease associations with altered neuronal developmental processes, abnormal emotional/affective behavior, mood disorders and MDD, among others, as being rescued by H3.3Q5A manipulations. Finally, given that both fluoxetine and H3.3Q5A mediated reductions of H3K4me3Q5ser in DRN were sufficient to reverse stress-induced dynamics of the mark and rescue stress-induced gene expression/behavior, we next sought to explore whether these two manipulations might induce alterations at the same genes, thereby linking fluoxetine’s genomic and behavioral rescue effects to the impact of directly manipulating the mark in the context of chronic stress. In doing we, we identified significant overlaps (odds ratio = 2.6; p = 3.6e-146) between PCGs displaying altered serotonylation dynamics in response to fluoxetine exposures in stress-susceptible mice and genes that were found to be differentially expressed following H3.3Q5A manipulations in the context of CSDS (**Fig. S6E**), with these overlapping genes displaying significant enrichment (FDR<0.05) for pathways/processes (GO Biological process) related to synaptic organization/function, as well as disease associated pathways (GWAS catalog) related to mood disorders (e.g., MDD, bipolar disorder, depressive symptoms) and other psychiatric illnesses (**Fig. S6F, Supplemental Data Tables 68-69**). In addition, we found that 37% of all PCGs exhibiting differential enrichment for H3K4me3Q5ser in human MDD cases + ADs *vs.* MDD subjects – ADs at their time of death overlapped with genes displaying significant differential expression between H3.3Q5A *vs.* empty CSDS mice, again indicating that reductions in H3K4me3Q5ser in DRN may contribute importantly to the regulation of genes associated with AD responsiveness. In sum, our viral manipulation data demonstrate that downregulation of H3K4me3Q5ser in DRN of chronically stressed mice is causally sufficient to reverse stress-mediated transcriptional programs and promote behavioral resilience. However, whether such downregulation of the mark following stress exposures (as opposed to prophylactic inhibition, as in the experiments presented above) would also be sufficient to ameliorate stress-induced deficits remains to be elucidated in future studies.

## DISCUSSION

Here, we demonstrated that DRN, the primary hub of serotonergic projection neurons in the central nervous system, displays robust transcriptional changes as a consequence of chronic social stress in both male and female mice. The biological processes predicted to be affected by chronic stress-related gene expression programs were found to be largely overlapping between the two sexes and significantly implicated disease associations with psychiatric and/or mood-related disorders, including MDD. These alterations in gene expression coincided with disruptions in H3 serotonylation dynamics in both male and female DRN, with similar results observed in postmortem tissues from individuals diagnosed with MDD. Interestingly, male mice deemed to be stress-resilient following CSDS displayed significant rescue of these H3K4me3Q5ser dynamics, indicating that patterns of differential H3K4me3Q5ser enrichment observed in stress-susceptible mice may contribute importantly to maladaptive behaviors elicited by chronic stress. We also observed in animals classified as being stress-susceptible (*vs.* stress-resilient) that the mark displayed aberrant accumulation in DRN during protracted periods following stress exposures and was largely rescued in response to chronic fluoxetine exposures, treatments that significantly reversed behavioral deficits observed in susceptible animals. Finally, we showed that directly reducing levels of H3 serotonylation in DRN prior to CSDS promoted behavioral resilience to chronic stress and significantly rescued stress-mediated gene expression programs, with many of the same genes displaying regulation by both chronic fluoxetine exposures and direct manipulations of H3Q5ser itself. In sum, these data establish a non-canonical, neurotransmission-independent role for 5-HT in the precipitation of stress-induced gene expression programs and maladaptive behavioral plasticity in DRN, results that suggest potential alternative roles for this important molecule in affect-related pathophysiology and the treatment of such disorders by classical SSRI ADs.

While the ‘5-HT hypothesis of depression’ remains highly influential, largely owing to the fact that most currently prescribed ADs act pharmacologically to increase 5-HT signaling in brain (as well in peripheral systems), a paucity of data exists directly implicating disruptions in serotonergic signaling/neurotransmission in the precipitation of disease. In fact, one recent meta-analysis attempting to link 5-HT (as well as the 5-HT metabolite 5-HIAA) concentrations in body fluids, serotonin 5-HT_1A_ receptor binding, SERT levels via imaging or at postmortem, tryptophan depletion studies or SERT gene-environment interactions to MDD pathology identified only weak, and often inconsistent evidence of interactions between these phenomena and MDD diagnosis in humans (5). Here, we posit that additional, previously undescribed 5-HT-related mechanisms may also contribute importantly to the pathophysiology of stress/mood-related disorders and should be considered in future studies aimed at examining functions for this molecule as a precipitating factor in disease.

Given that H3 serotonylation functions independently of neurotransmission and is critically important for both the establishment and maintenance of normal gene expression programs in brain, our observation that chronic stress, widely accepted as a major contributor to MDD pathology and incidence levels in humans, significantly alters baseline patterns of H3K4me3Q5ser in DRN – a phenomenon that if rescued (either through the use of viral vectors or chronic AD treatments), appears sufficient to restore stress-mediated gene expression programs and promote behavioral resilience – suggests that elementary correlations between 5-HT signaling (i.e., 5-HT levels and/or receptor binding) and MDD diagnosis may be insufficient to fully elucidate roles for this molecule in affect-related disorders. Furthermore, we hypothesize that these findings may help to explain, at least in part, the delayed efficacy of 5-HT associated ADs in both humans with MDD and preclinical rodent models. Many of our previous findings have suggested that H3 monoaminylation levels are largely dictated by intracellular donor (i.e., monoamine) concentrations (20), but once established in neuronal chromatin, it remains unclear how quickly the marks will be turned over, especially given the relatively slow kinetics of histone turnover observed in both neurons and glia (62). As such, AD treatments may function to increase 5-HT release from serotonergic neurons, thereby reducing intracellular 5-HT concentrations, eventually leading to loss, or restoration, of the mark within these cells. However, if the mark remains relatively stable during initial AD treatments, then its accumulation may not be fully resolved by acute administrations of these drugs. If true, then chronic treatments with ADs may be required to facilitate the full restoration of normal H3 serotonylation levels in serotonergic neurons, only with time would aberrant stress-induced gene expression programs be appropriately corrected. Further investigations will be needed to fully elucidate the precise kinetics of H3 serotonylation turnover in DRN as a consequence of AD treatments in order to demonstrate whether such dynamics are indeed causally linked to symptomatic alleviation of stress-related phenotypes. In addition, it also remains unknown how altering H3K4me3Q5ser levels in important 5-HTergic projection regions, such as mPFC, might affect phenotypic outcomes resulting from chronic stress and/or AD exposures. This will be important given that chronic fluoxetine exposures might be expected to simultaneously reduce H3K4me3Q5ser levels in DRN (which ameliorates stress-induced phenotypes) while increasing the mark in regions of the brain receiving 5-HTergic innervation. Such potential phenomena will require future studies to fully be resolved.

Additionally, while our current study is focused primarily on alterations in H3 serotonylation dynamics in DRN as a putative precipitating factor in stress-related gene expression programs and behavior, it is important to note that DRN is not a homogeneous monoaminergic brain structure, as it has been shown previously that a smaller population of dopaminergic neurons also reside in DRN and can contribute importantly to certain affect-related behaviors (63). This is of particular interest given that our previous work also identified dopamine as an important donor molecule for H3Q5 transamidation in brain, a modification (i.e., H3Q5dop) that we showed accumulates in ventral tegmental area (VTA) of rats during abstinence from chronic, volitional administration of cocaine and heroin (19,25). H3Q5dop accumulation was found to potentiate aberrant gene expression programs in VTA that contribute to hyper-dopamine release dynamics in response to drug cues and increased vulnerability to drug relapse-related behaviors (19). Like that of H3 serotonylation in DRN, which displayed acute downregulation following CSDS (24 hr after SI testing) and subsequent accumulation during protracted periods after chronic stress exposures, H3Q5dop was also found to be reduced in VTA immediately after drug administration, dynamics that were reversed during drug abstinence and were found to promote persistent maladaptive plasticity and increased cue-induced craving for drugs of abuse. Consistent with these earlier drug abuse studies, we found that the persistent accumulation of H3 serotonylation in DRN following chronic stress exposures influenced the potentiation of stress susceptibility. In addition, while at first glance, our data demonstrating that H3K4me3Q5ser levels (via western blotting) were reduced in individuals diagnosed with MDD (without ADs onboard at their time of death) may appear to contradict our mechanistic findings that H3 serotonylation accumulation in DRN is most tightly associated with stress-susceptibility, we posit that such reductions in human DRN are likely reflective of the agonal state of the subjects examined, as nearly all of the MDD –AD individuals included in this study died by suicide. Thus, it is possible that the molecular alterations in H3 serotonylation levels being captured in our data more closely resemble periods of ongoing stress, which would be consistent with our rodent data from 24 hr post-SI testing. Similar results were observed for H3Q5dop in VTA of postmortem subjects diagnosed with cocaine-dependence and who died by drug overdose, where we found that their global levels of H3Q5dop were downregulated and more closely resembled periods of active drug-taking in rodents (19). Thus, while comparisons of such molecular phenomena in preclinical rodent models *vs.* clinically diagnosed humans remain grossly informative, these types of postmortem human analyses may not faithfully inform on the precise mechanistic roles for H3 serotonylation in disease etiology, thereby further highlighting the importance of using well-controlled, preclinical models for the study of complex psychiatric disorders. It is important to note, however, that AD treatments in MDD patients were observed to renormalize total levels of H3K4me3Q5ser (via western blotting) in DRN with a greater number of PCGs displaying loss of the mark (as assessed via ChIP-seq), data that are consistent with the effects of chronic fluoxetine treatments observed in stress-susceptible animals. These findings indicate that alterations in the mark’s enrichment observed in behaviorally responsive, fluoxetine treated CSDS mice may indeed be of clinical relevance and may reflect functional chromatin adaptations that occur in human MDD subjects undergoing AD treatments.

An additional limitation of the current study is the possibility that our viral dominant negative approach may also impact H3Q5dop in DRN, as both marks are indeed present within this brain region (note that H3Q5his is only very weakly found within DRN (20)), although their relative stoichiometries remain unclear. Presumably, given that the proportion of serotonergic *vs.* dopaminergic neurons is largely skewed towards that of serotonergic cells in DRN, one might assume that the serotonylation mark would be more dominantly expressed, though this has yet to be tested empirically. While H3K4me3Q5ser and H3K4me3Q5dop are predicted to have similar molecular functions (e.g., recruiting the same “reader” proteins) it will be important in future studies to develop methodologies that can selectively target each modification independently (note that no such methodologies currently exist), followed by examinations of whether H3K4me3Q5dop (*vs.* H3K4me3Q5ser) is similarly responsive to chronic stress exposures in DRN. Further investigation of monoaminyl marks in other brain structures and cell populations beyond monoaminergic neurons may also uncover distinct regional or cell type-specific mechanisms that influence neuronal signaling and behavior. Finally, while histone H3 has been demonstrated to be a critical substrate for monoaminylation events in brain, future studies aimed at uncovering the full repertoire of monoaminylated proteins in brain, as well as their responsiveness to chronic stress exposures and AD treatments, may prove informative to the understanding of how alterations in monoaminergic activities may contribute to MDD pathophysiology and its treatment.

## ACKNOWLEDGEMENTS

We would like to thank members of the Maze and Russo laboratories for critical readings of the manuscript. This work was partially supported by grants from the National Institutes of Health: R01 MH116900 (I.M.), F99 NS125774 (S.L.F.), F31 MH116588 (S.L.F.), K99 MH120334 (L.A.F.), F32 MH125634 (E.L.N.), F32 MH126534 (J.C.C.), as well as funds from MQ (I.M.) Alfred P. Sloan Foundation (I.M.), One Mind (I.M.) and Howard Hughes Medical Institute (I.M.).

## CONTRIBUTIONS

A.A. and I.M. conceived of the project, designed the experiments, and interpreted the data; A.A., S.L.F., G.D.S., J.C.C., L.A.F., A.E.L., R.M.B., L.K., F.C., E.L.N., C.M., P.S., Y.L. and H.E.C. collected and analyzed the data; A.A., S.L.F., A.R., L.S. and I.M. performed the bioinformatics analyses; K.G. and C.A.T. provided human postmortem tissues; A.A. and I.M. wrote the manuscript.

## DATA & MATERIALS AVAILABILITY

The RNA-seq and ChIP-seq data generated in this study have been deposited in the National Center for Biotechnology Information Gene Expression Omnibus (GEO) database under accession number GSE216104. We declare that the data supporting findings for this study are available within the article and Supplementary Information. Related data are available from the corresponding author upon reasonable request. No restrictions on data availability apply.

## SUPPLEMENTAL MATERIALS

1) Title Page

2) Supplemental Data Figs. 1-7 with Captions

3) Captions for Supplemental Data Tables 1-71

**Other Supplementary Material for this manuscript includes the following:**

Supplemental Data Tables 1-71

## SUPPLEMENTAL MATERIALS

**Supplemental Data Fig. 1.**
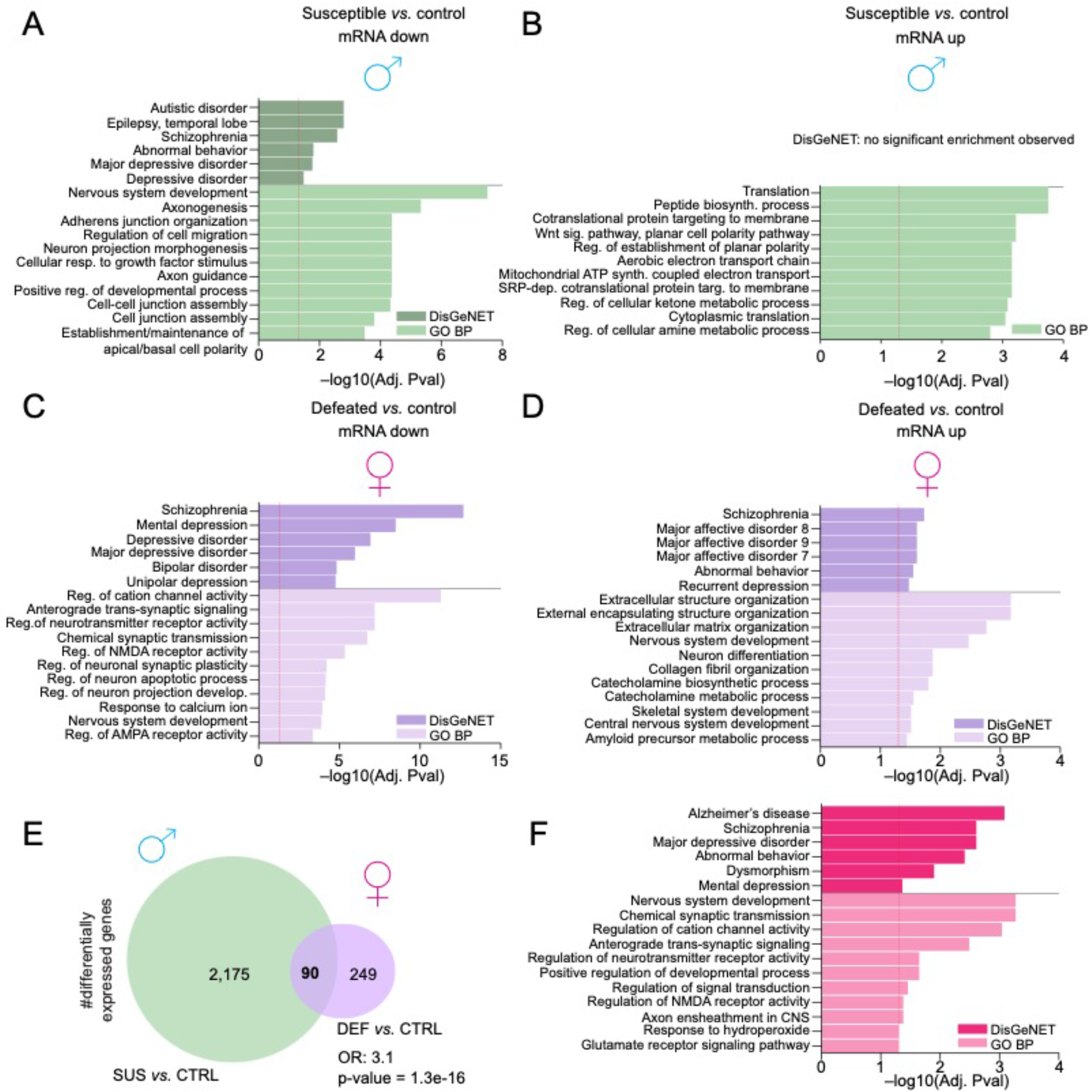
Directionality and overlap of gene expression changes observed following CSDS in males *vs.* females. **(A)** Example GO Biological Process and DisGeNET pathway enrichment (FDR<0.05) for the PCGs significantly downregulated (at FDR<0.1) in susceptible *vs.* control males. Dashed line indicates significance via adjusted p-value. **(B)** Example GO Biological Process and DisGeNET pathway enrichment (FDR<0.05) for the PCGs significantly upregulated (at FDR<0.1) in susceptible *vs.* control males. Dashed line indicates significance via adjusted p-value. **(C)** Example GO Biological Process and DisGeNET pathway enrichment (FDR<0.05) for the PCGs significantly downregulated (at FDR<0.1) in defeated *vs.* control females. Dashed line indicates significance via adjusted p-value. **(D)** Example GO Biological Process and DisGeNET pathway enrichment (FDR<0.05) for the PCGs significantly upregulated (at FDR<0.1) in defeated *vs.* control females. Dashed line indicates significance via adjusted p-value. **(E)** Venn diagram depicting the overlap (independent of directionality) between differentially expressed PCGs in male susceptible *vs.* control and female defeated *vs.* control. Odds ratio (OR) and respective p-value of overlap is provided. **(F)** Example GO Biological Process and DisGeNET pathway enrichment (FDR<0.05) for the differentially expressed PCGs significantly overlapping between male *vs.* female.

**Supplemental Data Fig. 2.**
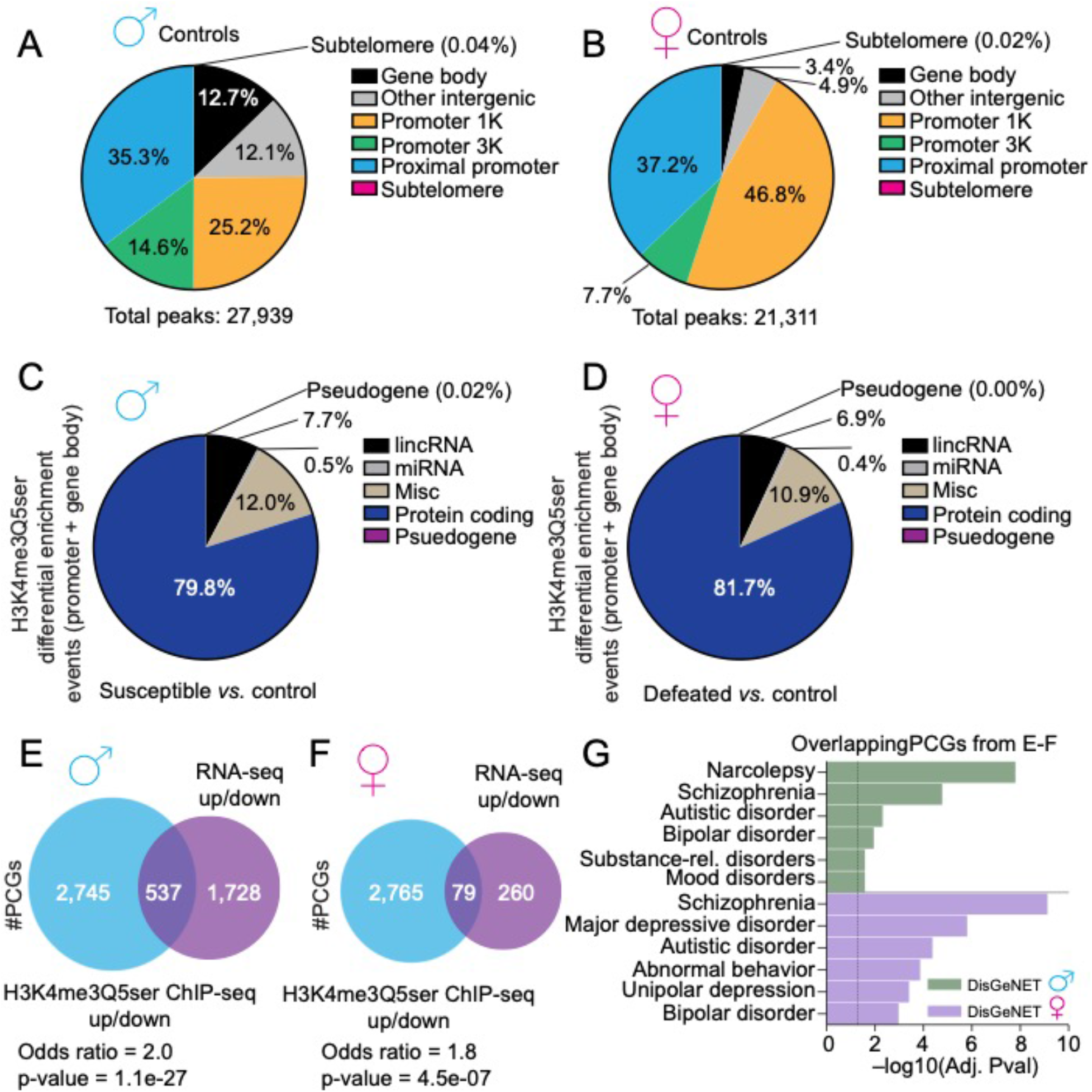
Genomic distributions of H3K4me3Q5ser baseline and differential enrichment, and correlations with differential gene expression. Pie charts describing the genomic distribution of H3K4me3Q5ser peaks (FDR<0.05, >5-fold enrichment over DNA input) in control **(A)** male *vs.* **(B)** female DRN. Pie charts describing the genomic distribution of H3K4me3Q5ser differential enrichment events (FDR<0.05, ≥ 1.5 or ≤ – 1.5) in promoters and gene bodies of **(C)** male susceptible *vs.* control or **(D)** female defeated *vs.* control DRN broken down by gene ‘type.’ Venn diagrams describing the overlap between differentially expressed PCGs (FDR<0.1, irrespective of directionality) and PCGs displaying differential H3K4me3Q5ser enrichment (FDR<0.05, ≥ 1.5 or ≤ –1.5 fold change, irrespective of directionality) in **(E)** male susceptible *vs.* control or **(F)** female defeated *vs.* control DRN. Odds ratios (OR) and respective p-values of overlap are provided. **(G)** Example DisGeNET pathway enrichment (FDR<0.05) for overlapping PCGs identified in E-F for males and females. Dashed line indicates significance via adjusted p-value.

**Supplemental Data Fig. 3.**
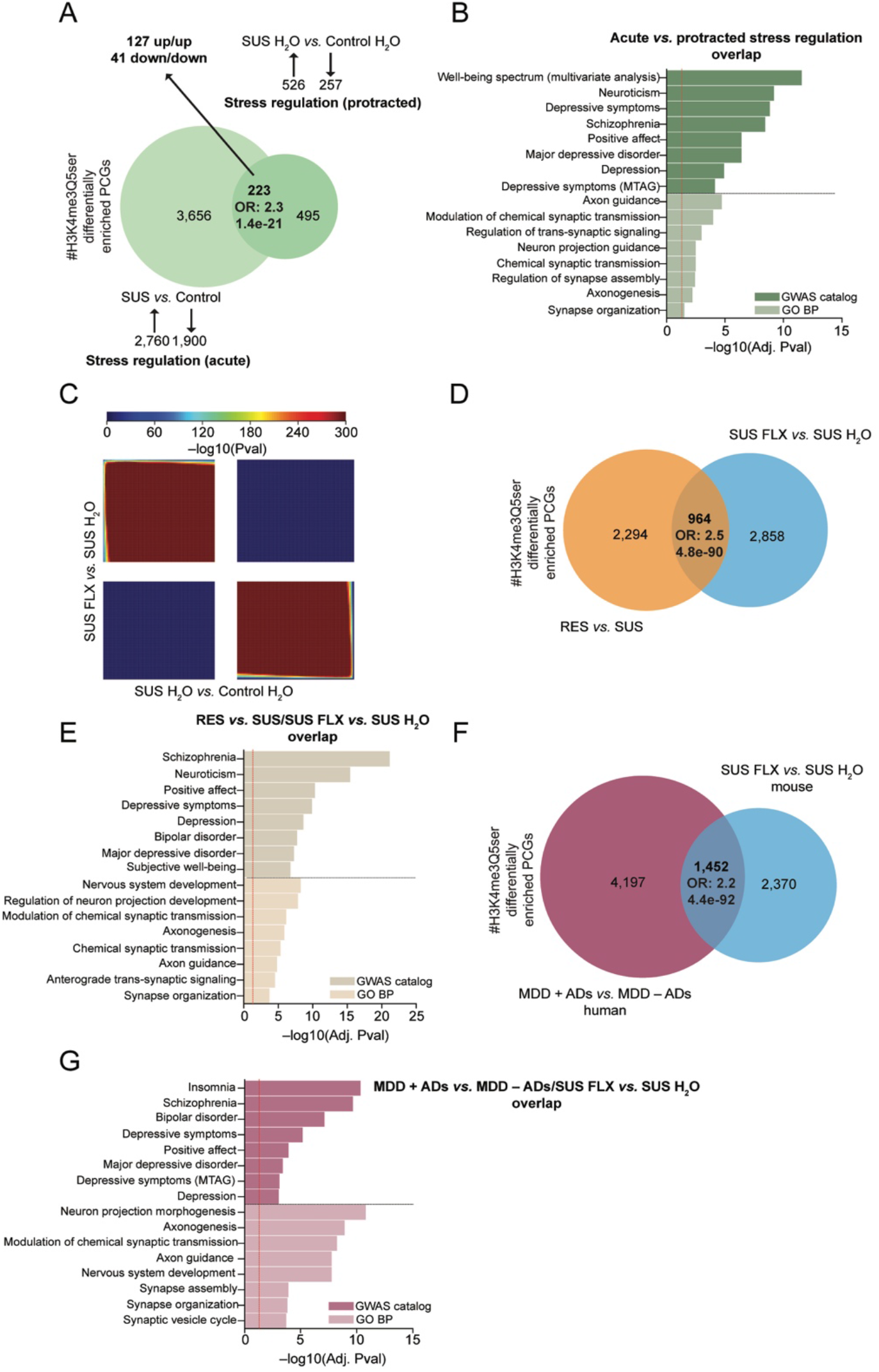
Chronic fluoxetine in CSDS male mice regulates H3K4me3Q5ser dynamics in DRN at PCGs relevant to stress-resilience and AD responsiveness in humans with MDD. **(A)** Venn diagram depicting the overlap (independent of directionality) between PCGs differentially enriched for H3K4me3Q5ser in male susceptible *vs.* control (24 hr post-SI testing; acute stress regulation) and male susceptible H_2_O *vs.* control H_2_O mice (30d post-initial SI testing; protracted stress regulation). Odds ratio (OR) and respective p-value of overlap is provided. **(B)** Example GO Biological Process and GWAS Catalog pathway enrichment (FDR<0.05) for the differentially enriched PCGs significantly overlapping between the two comparisons in A. **(C)** Threshold-free RRHO analyses comparing transcriptional profiles for stress-regulated genes in the absence or presence of fluoxetine at protracted periods following CSDS (*n* = 4-5/group), demonstrating that chronic fluoxetine treatments significantly reversed gene expression programs observed in response to stress in vehicle treated animals. Each pixel represents the overlap between differential transcriptomes, with the significance of overlap of a hypergeometric test color-coded. **(D)** Venn diagram depicting the overlap (independent of directionality) between PCGs differentially enriched for H3K4me3Q5ser in male resilient *vs.* stress-susceptible (24 hr post-SI testing) and male susceptible FLX *vs.* susceptible H_2_O mice (30d post-initial SI testing). Odds ratio (OR) and respective p-value of overlap is provided. **(E)** Example GO Biological Process and GWAS Catalog pathway enrichment (FDR<0.05) for the differentially enriched PCGs significantly overlapping between the two comparisons in D. **(F)** Venn diagram depicting the overlap (independent of directionality) between PCGs differentially enriched for H3K4me3Q5ser in male susceptible FLX *vs.* susceptible H_2_O (30d hr post-initial SI testing) and human MDD subjects −/+ ADs onboard at their time of death (*n* = 5/group). Odds ratio (OR) and respective p-value of overlap is provided. **(G)** Example GO Biological Process and GWAS Catalog pathway enrichment (FDR<0.05) for the differentially enriched PCGs significantly overlapping between the two comparisons in F.

**Supplemental Data Fig. 4.**
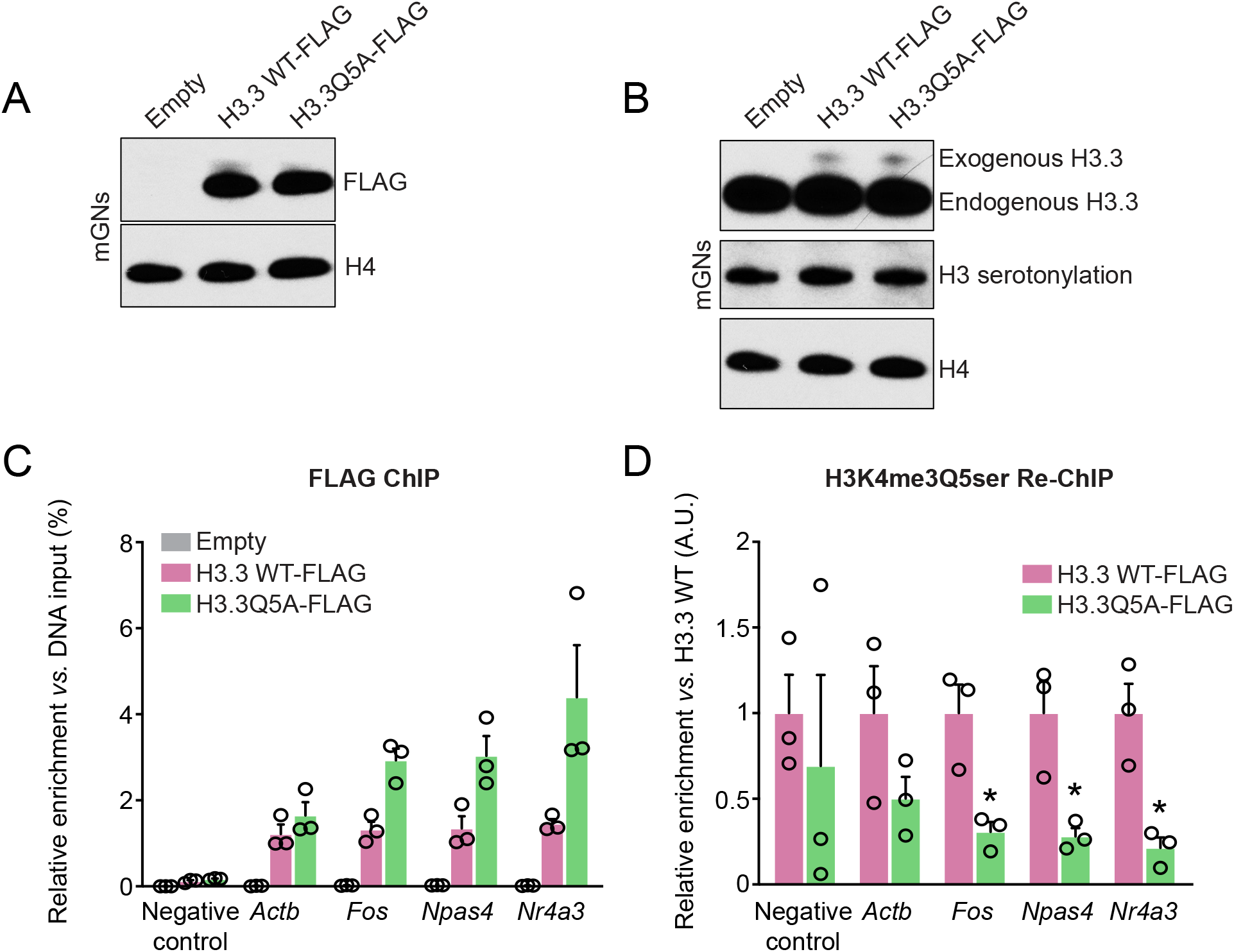
Validation of H3K4me3Q5ser downregulation in neurons at H3.3Q5A incorporated loci. **(A)** Western blotting validation of H3.3 WT-FLAG *vs.* H3.3Q5A-FLAG (*vs.* empty vector) expression – assessed using an anti-FLAG antibody – in cultured cerebellar granule neurons (cGNs). Total H4 was used as a loading control. **(B)** Western blotting-based comparison of endogenous *vs.* exogenous H3.3 expression (this antibody recognizes the globular domain of H3.3 and therefore can recognize both endogenous and exogenous H3.3) following transduction with empty *vs.* H3.3 WT-FLAG *vs.* H3.3Q5A-FLAG vectors in cGNs. Note that transduction with the aforementioned viruses (which account for ∼5% of the total H3.3 pool) did not result in a global downregulation of H3 serotonylation in these cells. Total H4 was used as a loading control. **(C)** FLAG ChIP-qPCRs following transduction with empty *vs.* H3.3 WT-FLAG *vs.* H3.3Q5A-FLAG vectors in cGNs demonstrating incorporation of exogenously expressed H3.3 (no signal observed in empty vector control expressing cells) proteins within the promoter of permissive genes but not within a negative control genomic locus. **(D)** FLAG ChIP-/H3K4me3Q5ser re-ChIP-qPCRs following transduction with empty *vs.* H3.3 WT-FLAG *vs.* H3.3Q5A-FLAG vectors in cGNs demonstrating significant loss of the serotonylation mark at H3.3Q5A *vs.* H3.3 WT incorporated genes, but not within a negative control locus (Student’s two-tailed t tests – Actb: *p*=0.1742, t_4_ = 1.650; *Fos*: *p*=0.0180, t_4_ = 3.939; *Npas4*: *p*=0.0210, t_4_ = 3.689; *Nr4a3*: *p*=0.0123, t_4_ = 4.335) empty vector ChIP-/re-ChIPs were excluded from this analysis based upon the lack of enrichment observed for this vector in C. Data presented as mean +/-SEM. A.U., arbitrary units; in D, data were normalized to respective enrichment of H3.3Q5A-FLAG *vs.* H3.3 WT-FLAG vectors to control for incorporation rates.

**Supplemental Data Fig. 5.**
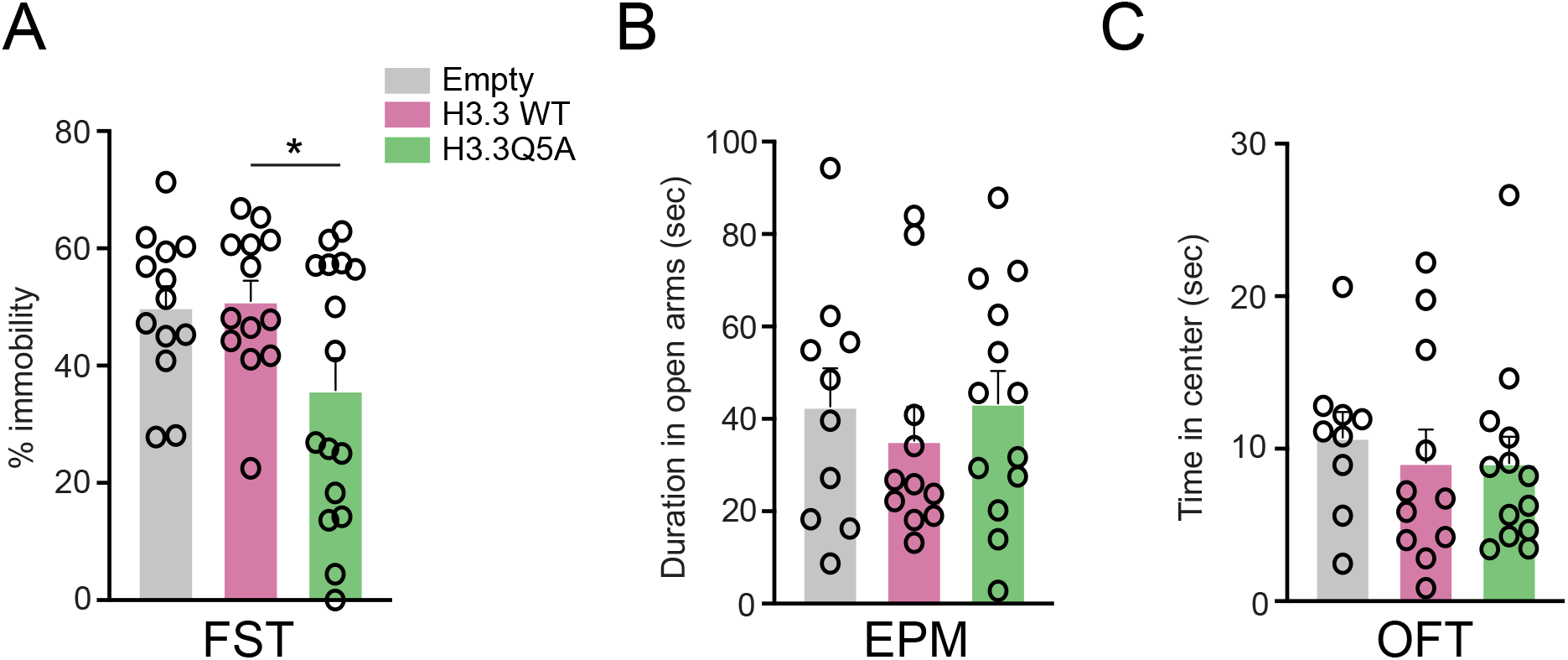
Reducing H3 serotonylation in DRN reduces behavioral despair in male mice, but does not affect anxiety-related behaviors. **(A)** Percent immobility in the forced swim test (FST) for control (i.e., non-stressed) male mice virally transduced intra-DRN with empty *vs.* H3.3WT *vs.* H3.3Q5A vectors (*n* = 13-16/group). Data were analyzed using a one-way ANOVA, with significant main effects (*p*=0.0319, F_2,39_ = 3.768). Tukey’s multiple comparisons test revealed significant differences between H3.3 WT *vs.* H3.3Q5A-transduced mice (*p*=0.05), with trending effects observed comparing empty *vs.* H3.3Q5A-transduced mice (*p*=0.07). **(B)** Duration of time spent in the open arms in the elevated plus maze (EPM) for control (i.e., non-stressed) male mice virally transduced intra-DRN with empty *vs.* H3.3WT *vs.* H3.3Q5A vectors (*n* = 10-13/group). Data were analyzed using a one-way ANOVA, with no significant main effects observed (*p*>0.05, F_2,31_ = 0.3605). **(C)** Duration of time spent in the center in the open field test (OFT) for control (i.e., non-stressed) male mice virally transduced intra-DRN with empty *vs.* H3.3WT *vs.* H3.3Q5A vectors (*n* = 9-13/group). Data were analyzed using a one-way ANOVA, with no significant main effects observed (*p*>0.05, F_2,30_ = 0.2260). Data presented as mean (+/-SEM).

**Supplemental Data Fig. 6.**
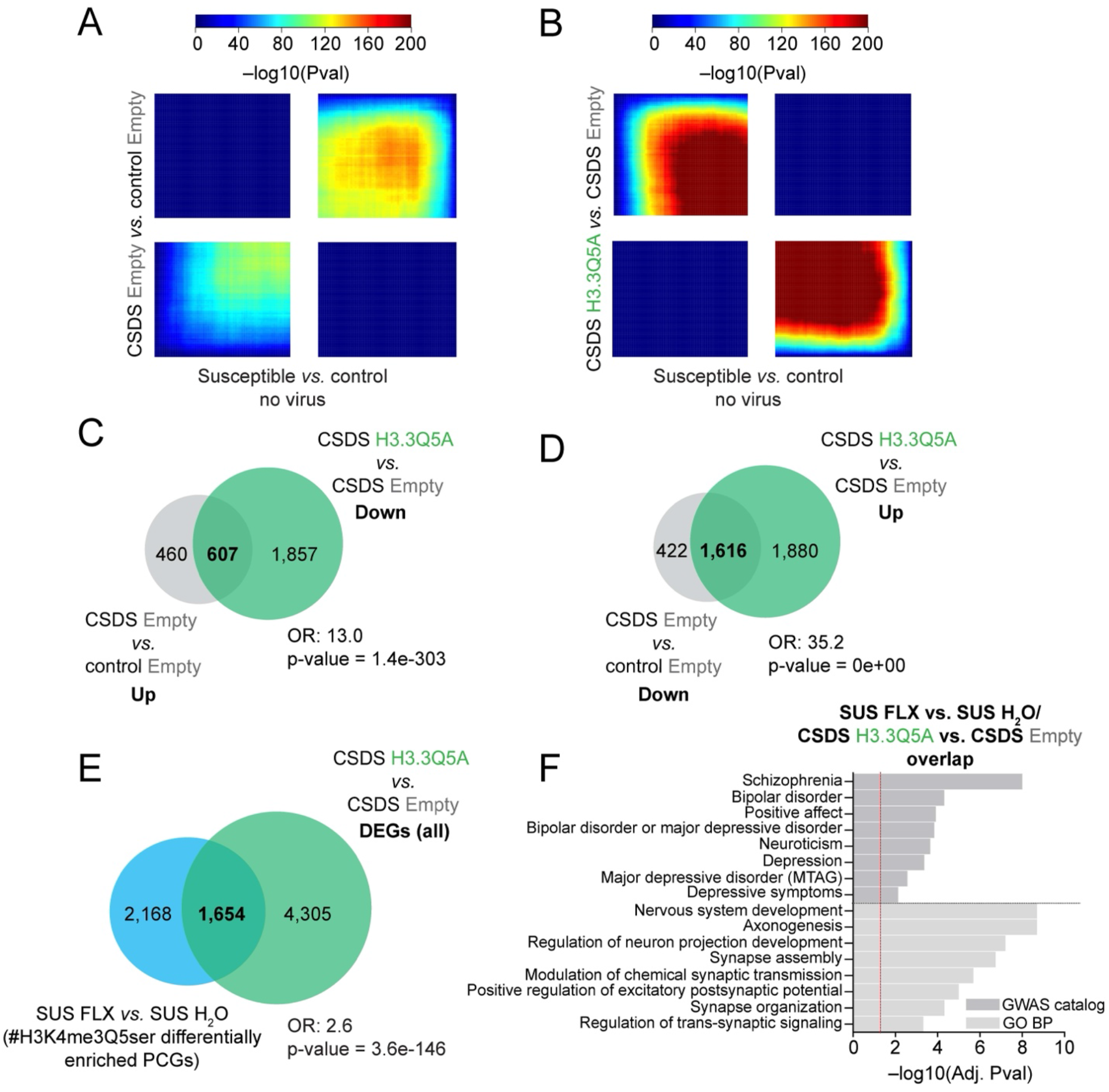
Correlations between gene expression profiles in DRN from susceptible *vs.* control male mice, −/+ viral transduction, and reversal of stress-induced gene expression in H3.3Q5A transduced animals. **(A)** Threshold-free RRHO analyses comparing transcriptional profiles for stress-regulated genes in empty vector (CSDS *vs.* control; *n* = 7-8/group) to non-virally transduced DRN from CSDS mice (*n* = 7-8/group). **(B)** Threshold-free RRHO analyses comparing transcriptional profiles for stress-regulated genes in susceptible *vs.* control (no virus; *n* = 7-8/group) male mice to H3.3Q5A CSDS *vs.* empty CSDS (*n* = 8-9/group). For A-B, each pixel represents the overlap between differential transcriptomes, with the significance of overlap of a hypergeometric test color-coded. Venn diagrams of overlap between PCGs displaying significant (**C)** upregulation (FDR<0.1) or **(D)** downregulation in their expression in CSDS empty *vs.* control empty comparisons and reversal of stress-induced gene expression following transduction with H3.3Q5A. Odds ratios (OR) and respective p-values of overlap are provided. **(E)** Venn diagram depicting the overlap (independent of directionality) between PCGs differentially enriched for H3K4me3Q5ser in male susceptible FLX *vs.* susceptible H_2_O mice (30d post-initial SI testing) and differentially expressed genes between H3.3Q5A CSDS *vs.* empty CSDS animals. Odds ratio (OR) and respective p-value of overlap is provided. **(F)** Example GO Biological Process and GWAS Catalog pathway enrichment (FDR<0.05) for the differentially enriched PCGs significantly overlapping between the two comparisons in E.

**Supplemental Data Fig. 7.**
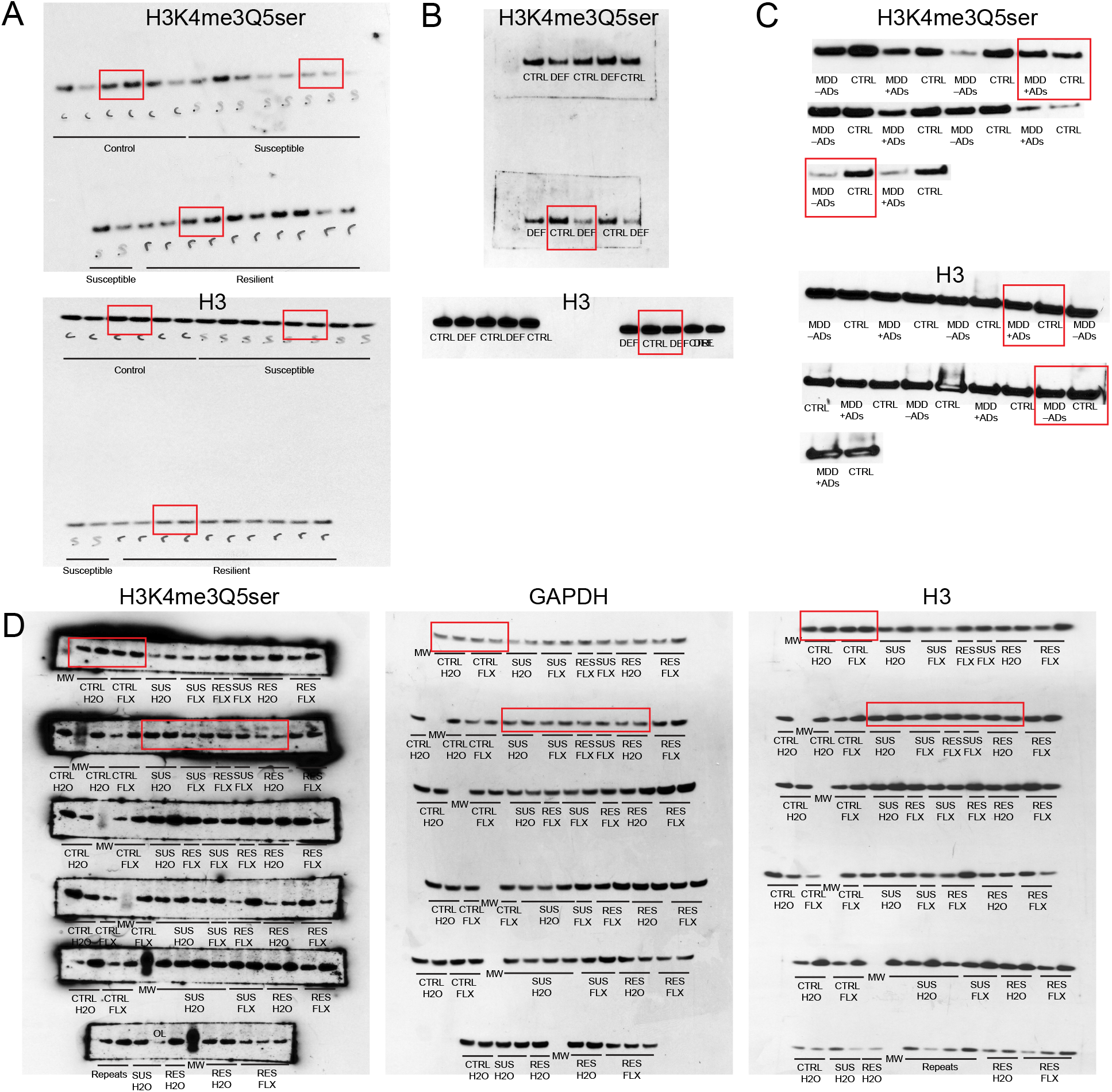
Uncropped western blots from Figures 2 and 3. Red rectangles indicate the representative blots displayed in the main Figures.

**SUPPLEMENTAL DATA TABLE CAPTIONS (with references to corresponding Figures)**

Excel file including Supplemental (Extended) Data Tables 1-71, which contain processed ChIP-seq and RNA-seq from adult male and female DRN, as well as human demographic information and mouse ChIP-qPCR primers.

**Supplemental Data Table 1:** Adult DRN (male) RNA-seq data (Susceptible vs. Control All), FDR<0.1 – Fig. 1B-C & Fig. S1A-B, E-F

**Supplemental Data Table 2:** Adult DRN (male) RNA-seq data (Susceptible vs. Control All), FDR<0.1, DisGeNET (FDR<0.05) – Fig. 1C

**Supplemental Data Table 3:** Adult DRN (male) RNA-seq data (Susceptible vs. Control All), FDR<0.1, Go BP (FDR<0.05) – Fig. 1C

**Supplemental Data Table 4:** Adult DRN (male) RNA-seq data (Susceptible vs. Control Down), FDR<0.1, DisGeNET (FDR<0.05) –Fig. S1A

**Supplemental Data Table 5:** Adult DRN (male) RNA-seq data (Susceptible vs. Control Up), FDR<0.1, DisGeNET (FDR<0.05) –Fig. S1B

**Supplemental Data Table 6:** Adult DRN (male) RNA-seq data (Susceptible vs. Control Down), FDR<0.1, GO BP (FDR<0.05) –Fig. S1A

**Supplemental Data Table 7:** Adult DRN (male) RNA-seq data (Susceptible vs. Control Up), FDR<0.1, GO BP (FDR<0.05) –Fig. S1B

**Supplemental Data Table 8:** Adult DRN (male) RNA-seq data (Susceptible vs. Resilient All), FDR<0.1 – Fig. 1B

**Supplemental Data Table 9:** Adult DRN (male) RNA-seq data (Resilient vs. Control All), FDR<0.1 – Fig. 1B

**Supplemental Data Table 10:** Adult DRN (female) RNA-seq data (Defeated vs. Control All), FDR<0.1 – Fig. 1E-F & Fig. S1C-F

**Supplemental Data Table 11:** Adult DRN (female) RNA-seq data (Defeated vs. Control All), FDR<0.1, DisGeNET (FDR<0.05) – Fig. 1F

**Supplemental Data Table 12:** Adult DRN (female) RNA-seq data (Defeated vs. Control All), FDR<0.1, Go BP (FDR<0.05) – Fig. 1F

**Supplemental Data Table 13:** Adult DRN (female) RNA-seq data (Defeated vs. Control Down), FDR<0.1, DisGeNET (FDR<0.05) –Fig. S1C

**Supplemental Data Table 14:** Adult DRN (female) RNA-seq data (Defeated vs. Control Up), FDR<0.1, DisGeNET (FDR<0.05) –Fig. S1D

**Supplemental Data Table 15:** Adult DRN (female) RNA-seq data (Defeated vs. Control Down), FDR<0.1, GO BP (FDR<0.05) –Fig. S1C

**Supplemental Data Table 16:** Adult DRN (female) RNA-seq data (Defeated vs. Control Up), FDR<0.1, GO BP (FDR<0.05) –Fig. S1D

**Supplemental Data Table 17:** Adult DRN RNA-seq data (Male Susceptible vs. Control & Female Defeated vs. Control Overlap All), FDR<0.1, DisGeNET (FDR<0.05) –Fig. S1F

**Supplemental Data Table 18:** Adult DRN RNA-seq data (Male Susceptible vs. Control & Female Defeated vs. Control Overlap All), FDR<0.1, Go BP (FDR<0.05) –Fig. S1F

**Supplemental Data Table 19:** Adult DRN (male) H3K4me3Q5ser ChIP-seq peaks (Control), FDR<0.05, >5-fold enrichment over input – Figure 2D-I & Fig. S2A-B

**Supplemental Data Table 20:** Adult DRN (male) H3K4me3Q5ser ChIP-seq peaks (Susceptible), FDR<0.05, >5-fold enrichment over input – Figure 2D-I & Fig. S2A-B

**Supplemental Data Table 21:** Adult DRN (male) H3K4me3Q5ser ChIP-seq peaks (Resilient), FDR<0.05, >5-fold enrichment over input – Figure 2D-I & Fig. S2A-B

**Supplemental Data Table 22:** Adult DRN (female) H3K4me3Q5ser ChIP-seq peaks (Control), FDR<0.05, >5-fold enrichment over input – Figure 2D-G & Fig. S2A-B

**Supplemental Data Table 23:** Adult DRN (female) H3K4me3Q5ser ChIP-seq peaks (Defeated), FDR<0.05, >5-fold enrichment over input – Figure 2D-G & Fig. S2A-B

**Supplemental Data Table 24:** Adult DRN (male) H3K4me3Q5ser diffReps (Susceptible vs. Control All, FDR<0.05, log2FC ≥ 1.5 or ≤ –1.5 – Fig. 2F-G & Fig. S2C, E, G

**Supplemental Data Table 25:** Adult DRN (male) H3K4me3Q5ser diffReps (Susceptible vs. Resilient All, FDR<0.05, log2FC ≥ 1.5 or ≤ –1.5 – Fig. 2H-I & Fig. S2C, E, G

**Supplemental Data Table 26:** Adult DRN (male) H3K4me3Q5ser diffReps (Resilient vs. Control All, FDR<0.05, log2FC ≥ 1.5 or ≤ –1.5 – Fig. 2

**Supplemental Data Table 27:** Adult DRN (female) H3K4me3Q5ser diffReps (Defeated vs. Control All, FDR<0.05, log2FC ≥ 1.5 or ≤ –1.5 – Fig. 2F-G & Fig. S2D, F, G

**Supplemental Data Table 28:** Adult DRN ChIP-seq data (Male Susceptible vs. Control & Female Defeated vs. Control Overlap All), FDR<0.05, log2FC ≥ 1.5 or ≤ –1.5, GWAS Catalog (FDR<0.05) – Fig. 2G

**Supplemental Data Table 29:** Adult DRN ChIP-seq data (Male Susceptible vs. Control & Female Defeated vs. Control Overlap All), FDR<0.05, log2FC ≥ 1.5 or ≤ –1.5, DisGeNET (FDR<0.05) – Fig. 2G

**Supplemental Data Table 30:** Adult DRN ChIP-seq data (Male Susceptible vs. Control & Female Defeated vs. Control Overlap All), FDR<0.05, log2FC ≥ 1.5 or ≤ –1.5, GO BP (FDR<0.05) – Fig. 2G

**Supplemental Data Table 31:** Adult DRN ChIP-seq data (Male Susceptible vs. Control & Male Resilient vs. Susceptible Overlap, Reversed in Resilient), FDR<0.05, log2FC ≥ 1.5 or ≤ –1.5, GWAS Catalog (FDR<0.05) – Fig. 2I

**Supplemental Data Table 32:** Adult DRN ChIP-seq data (Male Susceptible vs. Control & Male Resilient vs. Susceptible Overlap, Reversed in Resilient), FDR<0.05, log2FC ≥ 1.5 or ≤ –1.5, DisGeNET (FDR<0.05) – Fig. 2I

**Supplemental Data Table 33:** Adult DRN ChIP-seq data (Male Susceptible vs. Control & Male Resilient vs. Susceptible Overlap, Reversed in Resilient), FDR<0.05, log2FC ≥ 1.5 or ≤ –1.5, GO BP (FDR<0.05) – Fig. 2I

**Supplemental Data Table 34:** Adult DRN ChIP-seq data (Male Susceptible vs. Control All; FDR<0.05, log2FC ≥ 1.5 or ≤ –1.5) vs. Adult DRN RNA-seq data (Male Susceptible vs. Control All; FDR<0.1), DisGeNET (FDR<0.05) –Fig. S2G (Overlapping PCGs)

**Supplemental Data Table 35:** Adult DRN ChIP-seq data (Female Defeated vs. Control All; FDR<0.05, log2FC ≥ 1.5 or ≤ –1.5) vs. Adult DRN RNA-seq data (Female Defeated vs. Control All; FDR<0.1), DisGeNET (FDR<0.05) –Fig. S2G (Overlapping PCGs)

**Supplemental Data Table 36:** Adult DRN (male) H3K4me3Q5ser ChIP-seq peaks (Control H2O), FDR<0.05, >5-fold enrichment over input – Figure 3E-G & Fig. S3A-B

**Supplemental Data Table 37:** Adult DRN (male) H3K4me3Q5ser ChIP-seq peaks (Susceptible H2O), FDR<0.05, >5-fold enrichment over input – Figure 3E-H & Fig. S3A-B, D-E

**Supplemental Data Table 38:** Adult DRN (male) H3K4me3Q5ser diffReps (Susceptible H2O vs. Control H2O All, FDR<0.05, log2FC ≥ 1.0 or ≤ –1.0 – Fig. 3E-G & Fig. S3A-B

**Supplemental Data Table 39:** Adult DRN (male) H3K4me3Q5ser diffReps (Susceptible vs. Control All, FDR<0.05, log2FC ≥ 1.0 or ≤ –1.0 – Fig. 3E-G & Fig. S3A-B

**Supplemental Data Table 40:** Adult DRN ChIP-seq data (Male Susceptible vs. Control Acute & Male Susceptible vs. Control Protracted Overlap All), FDR<0.05, log2FC ≥ 1.0 or ≤ –1.0, GO BP (FDR<0.05) –Fig. S3B

**Supplemental Data Table 41:** Adult DRN ChIP-seq data (Male Susceptible vs. Control Acute & Male Susceptible vs. Control Protracted Overlap All), FDR<0.05, log2FC ≥ 1.0 or ≤ –1.0, GWAS Catalog (FDR<0.05) –Fig. S3B

**Supplemental Data Table 42:** Adult DRN (male) RNA-seq data (Susceptible H2O vs. Control H2O All), –Fig. S3C

**Supplemental Data Table 43:** Adult DRN (male) RNA-seq data (Susceptible FLX vs. Control H2O All), –Fig. S3C

**Supplemental Data Table 44:** Adult DRN (male) H3K4me3Q5ser ChIP-seq peaks (Control FLX), FDR<0.05, >5-fold enrichment over input – Figure 3E-F

**Supplemental Data Table 45:** Adult DRN (male) H3K4me3Q5ser ChIP-seq peaks (Susceptible FLX), FDR<0.05, >5-fold enrichment over input – Figure 3E-H & Fig. S3D-G

**Supplemental Data Table 46:** Adult DRN (male) H3K4me3Q5ser diffReps (Susceptible FLX vs. Susceptible H2OAll, FDR<0.05, log2FC ≥ 1.0 or ≤ –1.0 – Fig. 3E-H & Fig. S3D-G

**Supplemental Data Table 47:** Adult DRN (male) H3K4me3Q5ser diffReps (Control FLX vs. Control H2O All, FDR<0.05, log2FC ≥ 1.0 or ≤ –1.0 – Fig. 3E-F

**Supplemental Data Table 48:** Adult (Male) DRN ChIP-seq data (Susceptible FLX vs. Susceptible H2O All), FDR<0.05, log2FC ≥ 1.0 or ≤ –1.0, GO BP (FDR<0.05) – Fig. 3H

**Supplemental Data Table 49:** Adult (Male) DRN ChIP-seq data (Susceptible FLX vs. Susceptible H2O All), FDR<0.05, log2FC ≥ 1.0 or ≤ –1.0, GWAS Catalog (FDR<0.05) – Fig. 3H

**Supplemental Data Table 50:** Adult DRN (male) H3K4me3Q5ser diffReps (Susceptible vs. Resilient All, FDR<0.05, log2FC ≥ 1.0 or ≤ –1.0 –Fig. S3D-E

**Supplemental Data Table 51:** Adult (Male) DRN ChIP-seq data (Susceptible FLX vs. Susceptible H2O & Resilient vs. Susceptible Overlap All), FDR<0.05, log2FC ≥ 1.0 or ≤ –1.0, GO BP (FDR<0.05) –Fig. S3E

**Supplemental Data Table 52:** Adult (Male) DRN ChIP-seq data (Susceptible FLX vs. Susceptible H2O & Resilient vs. Susceptible Overlap All), FDR<0.05, log2FC ≥ 1.0 or ≤ –1.0, GWAS Catalog (FDR<0.05) –Fig. S3E

**Supplemental Data Table 53:** Human DRN H3K4me3Q5ser ChIP-seq peaks (Matched Controls for MDD – ADs), FDR<0.05, >5-fold enrichment over input –Fig. S3F-G

**Supplemental Data Table 54:** Human DRN H3K4me3Q5ser ChIP-seq peaks (Matched Controls for MDD + ADs), FDR<0.05, >5-fold enrichment over input –Fig. S3F-G

**Supplemental Data Table 55:** Human DRN H3K4me3Q5ser ChIP-seq peaks (MDD – ADs), FDR<0.05, >5-fold enrichment over input –Fig. S3F-G

**Supplemental Data Table 56:** Human DRN H3K4me3Q5ser ChIP-seq peaks (MDD + ADs), FDR<0.05, >5-fold enrichment over input –Fig. S3F-G

**Supplemental Data Table 57:** Human DRN H3K4me3Q5ser diffReps (MDD – ADs vs. Matched Controls All, FDR<0.05, log2FC ≥ 1.0 or ≤ –1.0 –Fig. S3F-G

**Supplemental Data Table 58:** Human DRN H3K4me3Q5ser diffReps (MDD + ADs vs. MDD – ADs, FDR<0.05, log2FC ≥ 1.0 or ≤ –1.0 –Fig. S3F-G

**Supplemental Data Table 59:** Human DRN ChIP-seq data (MDD + ADs vs. MDD – ADs) vs. Mouse DRN (Male) ChIP-seq Data (Susceptible FLX vs. Susceptible H2O) Overlap All, FDR<0.05, log2FC ≥ 1.0 or ≤ –1.0, GO BP (FDR<0.05) –Fig. S3G

**Supplemental Data Table 60:** Human DRN ChIP-seq data (MDD + ADs vs. MDD – ADs) vs. Mouse DRN (Male) ChIP-seq Data (Susceptible FLX vs. Susceptible H2O) Overlap All, FDR<0.05, log2FC ≥ 1.0 or ≤ –1.0, GWAS Catalog (FDR<0.05) –Fig. S3G

**Supplemental Data Table 61:** Adult Viral DRN RNA-seq data (Male Empty CSDS vs. Empty Control All; FDR<0.1) – Fig. 4D-E & Fig. S5A-F

**Supplemental Data Table 62:** Adult Viral DRN RNA-seq data (Male H3.3 WT CSDS vs. H3.3 WT Control All; FDR<0.1) – Fig. 4D

**Supplemental Data Table 63:** Adult Viral DRN RNA-seq data (Male H3.3Q5A CSDS vs. Empty CSDS Control All; FDR<0.1) – Fig. 4D & Fig. S6B-F

**Supplemental Data Table 64:** Adult Viral DRN RNA-seq data (Male H3.3Q5A CSDS vs. H3.3 WT CSDS Control All; FDR<0.1) – Fig. 4D

**Supplemental Data Table 65:** Adult Viral DRN RNA-seq data (Male Empty CSDS vs. Empty Control & H3.3Q5A CSDS vs. Empty CSDS Control Reversed; FDR<0.1), GWAS Catalog (FDR<0.05) – Fig. 4E

**Supplemental Data Table 66:** Adult Viral DRN RNA-seq data (Male Empty CSDS vs. Empty Control & H3.3Q5A CSDS vs. Empty CSDS Control Reversed; FDR<0.1), DisGeNET (FDR<0.05) – Fig. 4E

**Supplemental Data Table 67:** Adult Viral DRN RNA-seq data (Male Empty CSDS vs. Empty Control & H3.3Q5A CSDS vs. Empty CSDS Control Reversed; FDR<0.1), GO BP (FDR<0.05) – Fig. 4E

**Supplemental Data Table 68:** Adult Viral DRN RNA-seq data (Male H3.3Q5A CSDS vs. Empty CSDS) and Adult DRN (Susceptible FLX vs. Susceptible H2O) Overlap All, FDR<0.1, GO BP (FDR<0.05) –Fig. S6F

**Supplemental Data Table 69:** Adult Viral DRN RNA-seq data (Male H3.3Q5A CSDS vs. Empty CSDS) and Adult DRN (Susceptible FLX vs. Susceptible H2O) Overlap All, FDR<0.1, GWAS Catalog (FDR<0.05) –Fig. S6F

**Supplemental Data Table 70:** Human postmortem DRN demographic information - Fig. 2C & Fig. S3F-G

**Supplemental Data Table 71:** Mouse qChIP Primers - Fig. S4

